# Neuronal differentiation affects tissue mechanics and progenitor arrangement in the vertebrate neuroepithelium

**DOI:** 10.1101/536342

**Authors:** Pilar Guerrero, Ruben Perez-Carrasco, Marcin Zagorski, David Page, Anna Kicheva, James Briscoe, Karen M. Page

## Abstract

Cell division, movement and differentiation contribute to pattern formation in developing tissues. This is the case in the vertebrate neural tube where neurons differentiate in a characteristic pattern from a highly dynamic proliferating pseudostratified epithelium. To investigate how progenitor proliferation and differentiation affect cell arrangement and growth of the neural tube, we use experimental measurements to develop a mechanical model of the apical surface of the neuroepithelium that incorporates inter-kinetic nuclear movement and spatially varying rates of neuronal differentiation. Simulations predict that tissue growth and the shape of lineage-related clones of cells differ with the rate of differentiation. Growth is isotropic in regions of high differentiation, but dorsoventrally biased in regions of low differentiation. This is consistent with experimental observations. The absence of directional signalling in the simulations indicates that global mechanical constraints are sufficient to explain the observed differences in anisotropy. This provides insight into how the tissue growth rate affects cell dynamics and growth anisotropy and opens up possibilities to study the coupling between mechanics, pattern formation and growth in the neural tube.

## INTRODUCTION

The mechanisms that control the arrangement of cells in developing tissues involve both molecular and mechanical processes that spatially and temporally coordinate the division, shape, displacement and differentiation of cells. A central challenge is to understand the interplay between tissue growth, pattern formation and the mechanical forces that act to shape tissues during development.

Studies of several systems have begun to provide insight into how these processes are coordinated (Alt et al. (2017); Merkel and Manning (2017)). For example, in the Drosophila wing imaginal disc a combination of experimental observations, quantitative image analysis, and computational modelling have revealed the global patterns of mechanical tension that affect the final size and shape of the wing. These patterns result from spatial differences in proliferation, cell shape and division orientation (Shraiman (2005); Kursawe et al. (2015); Aigouy et al. (2010); Campinho et al. (2013); LeGoff et al. (2013); Mao et al. (2013); Guirao et al. (2015); Aegerter-Wilmsen et al. (2010); Dye et al. (2017)), as well as external mechanical constraints, such as the attachment of the wing blade to the contracting wing hinge (Aigouy et al. (2010); Etournay et al. (2015); Ray et al. (2015)). Molecularly, wing morphogenesis is influenced by planar-polarity signalling, which influences the apical geometry of cells and the orientation of cell division (Mao et al. (2011); Aigouy et al. (2010)).

Similar to imaginal discs, the vertebrate neural tube is a pseudostratified epithelium. During neurulation the neuroepithelium folds at the ventral midline and closes dorsally to form a cylindrical neural tube, with the apical surfaces of neural progenitors facing the interior lumen (Gilbert (2014)). The proliferation of neural progenitors contributes to growth of the neural tube along the anterioposterior and dorsoventral axes. Additionally, proliferating cells undergo inter-kinetic nuclear movement (IKNM) during which each cell’s nucleus translocates along the apicobasal axis in synchrony with cell cycle progression (Sauer (1935)). A direct consequence of IKNM is that the apicobasal shape, the apical surface of cells, and the interactions between neighbouring cells change in a highly dynamic manner (reviewed in (Strzyz et al. (2016))).

At the same time as the neural tube grows, long range signals control patterning by regulating the expression of transcription factors within the tissue (reviewed in (Sagner et al. (2018))). The dynamics of this regulatory network results in the specification of molecularly distinct domains of progenitor subtypes arranged along the dorsoventral (DV) axis. The set of transcription factors expressed in a progenitor domain determines the subtype identity of neurons it generates. As neurons are formed, they delaminate from the epithelium to take up residence basally in the forming mantle zone. The delamination of newly born neurons contributes to the morphodynamics of the neuroepithelium, further reshaping cell-cell interactions and the arrangement of cells within the neural tube.

Previous studies of the neural tube indicated that patterning and growth are tightly coordinated. Cell death is negligible and the rate of progenitor proliferation is spatially uniform throughout the epithelium (Kicheva et al. (2014)). However, the rates of terminal neuronal differentiation vary depending on progenitor identity. Most notably, starting at mouse embryonic day E9.5, motor neuron progenitors (pMN) differentiate at a significantly faster rate than other progenitor subtypes (Kicheva et al. (2014); Ericson et al. (1996)). This difference in the rates of terminal differentiation correlated with a difference in clone shape in lineage tracing experiments (reproduced in Fig. 1A from (Kicheva et al. (2014))). In particular, while the spread of clones in all domains was similar, their DV spread was not. Clones in all but the pMN domain are elongated along the DV axis compared to the AP axis with an average ratio of AP to DV spread of ~ 0.3. By contrast, clones in the pMN domain have an average AP/DV ratio of ~ 1 indicating equal growth in DV and AP directions. Thus the higher differentiation rate of MN progenitors correlates with a specific change in the DV growth of clones. This raises the question of what mechanisms operate to ensure equivalent AP growth across the tissue, while at the same time allowing for cell-type specific differences in DV growth rates.

**Fig. 1.**
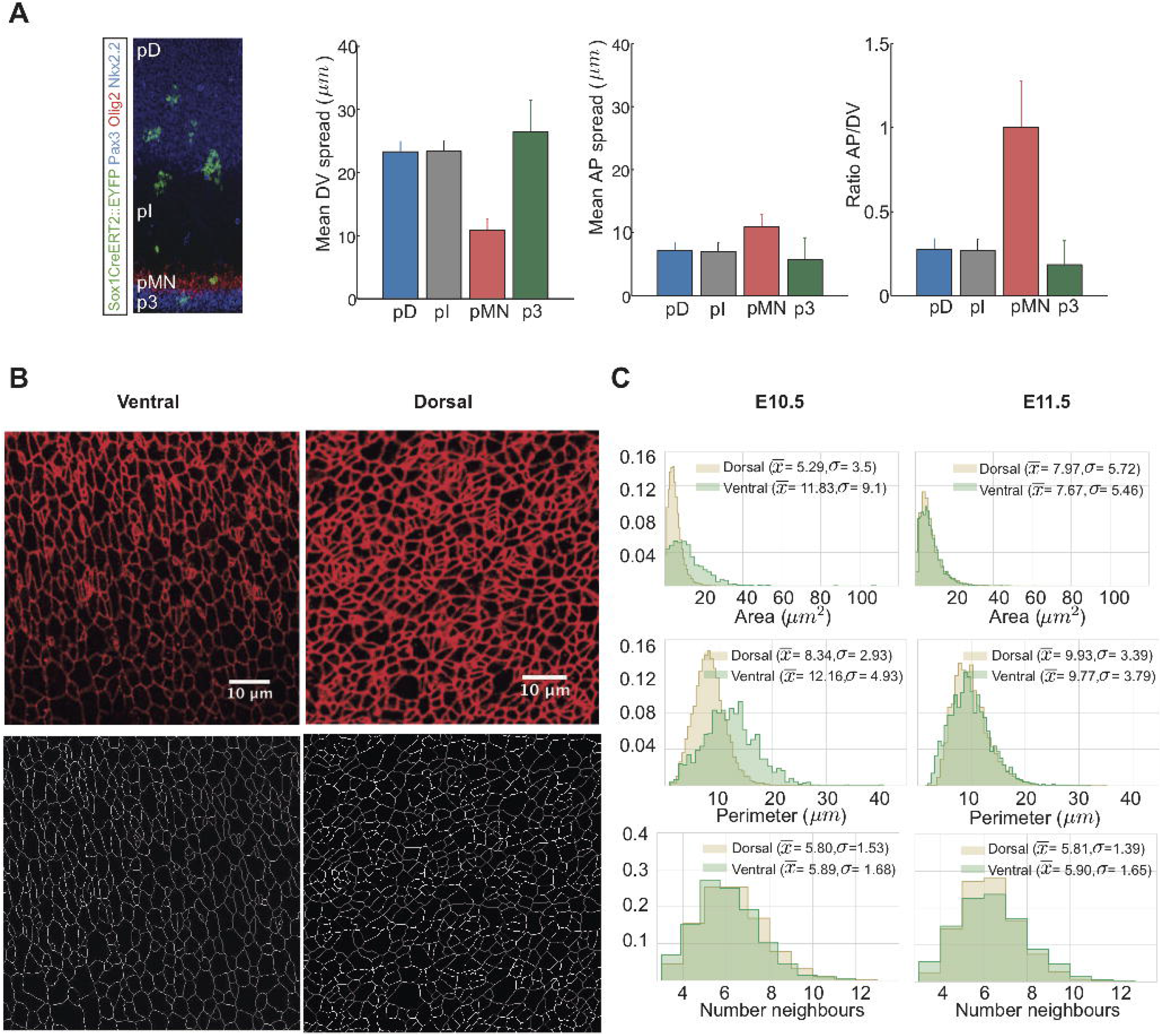
**A**) Analysis of clonal shape in E11.5 embryos, data from (Kicheva et al. (2014)). Clonal labelling was induced at E9.5 of development. The spread in AP direction is similar in all domains of the neural tube, but the DV spread, as well as the AP/DV ratio is different in the pMN domain. **B**) Top: Apical surface of E11.5 flat mounted mouse neural tube immunostained for ZO-1. Images were taken within the ventral and dorsal halves of the neural tube as indicated. Dorsal side up. Scale bar = 10 μm. Bottom: Segmented images after manual correction. **C**) Histograms of apical area, perimeter, and number of neighbours of cells from the dorsal (brown) and ventral (green) regions of E10.5 and E11.5 neural tubes. Sample sizes: E10.5, n=25 images of dorsal and 5 images of ventral domains from 7 different embryos;E11.5, 11 images of dorsal, 3 images of ventral domains from 3 embryos.

To address this, we developed computational tools to simulate the growth of the neuroepithelium and investigate the role of different mechanisms in the morphodynamics of the tissue. We made use of a representation of the apical 2D surface of the epithelium by employing a vertex model formalism (Nagai and Honda (2001); Fletcher et al. (2013); Asgari-Targhi (2012); Smith et al. (2011); Chiou et al. (2012); Canela-Xandri et al. (2011); Bock et al. (2010); Farhadifar et al. (2007)). Vertex models have been used successfully to describe mechanical and molecular influences that determine the tissue growth and form of several epithelia (Landsberg et al. (2009); Farhadifar et al. (2007); Aegerter-Wilmsen et al. (2010); Wartlick et al. (2011); Trichas et al. (2012); Salbreux et al. (2012)). In these models each cell is represented as a polygon, the vertices and edges of which are shared between adjacent cells. The dynamics of a cell are described by the movement of its vertices, which are controlled by adhesive/tensile, contractile and repelling forces in and between cells.

To take account of the 3D configuration of the neural tube, we incorporated the effects of IKNM into the simulation framework. Using experimental data from the mouse neural tube, we then established model parameters for which simulations match *in vivo* observations. We used the resulting model to explore the anisotropies of clonal shape within the neuroepithelium and the effect of incorporating spatially varying differentiation rates within the tissue. Strikingly, we found that the increased differentiation rate of pMN progenitors is sufficient to explain the different shape of clones within the pMN domain. This indicates that the differences in clonal shape arise from differences in progenitor differentiation rates and global mechanical constraints, and do not require polarised molecular signalling mechanisms. More generally, the availability of computational tools that accurately simulate the developing neuroepithelium will contribute to our understanding of how tissue patterning and growth are controlled and coordinated.

## RESULTS

### Cell geometry in the mouse neuroepithelium

To construct a mechanical model of neural tube growth we first measured key features of neural progenitor organisation in the mouse embryonic neural tube. To this end, we imaged the apical surface of the neural tube at forelimb level of E10.5 and E11.5 mouse embryos stained with ZO-1 to reveal tight junctions (Fig. 1B, top). The images were segmented and vertices and edges defined using ‘Packing Analyzer v2.0’ (Aigouy et al. (2010)) (Methods, Fig. 1B, bottom).

We collected images of the dorsal and ventral halves of the neural tube at E10.5 and E11.5. The dorsal images comprise the progenitors of dorsal interneuron subtypes, and we refer to this region as the pD domain. The ventral images cover motor neuron and intermediate neuron progenitor subtypes and we term this the pMN region (Methods). From the resulting segmented images we determined the distributions of cell areas, cell perimeters and number of neighbours per cell (Fig. 1C). Cells in all samples had on averaged 6 neighbours as expected (Gibson et al. (2006); Classen et al. (2005)), with a standard deviation of ~1.5. There were some differences in the mean and variance of cell areas and perimeters in the samples (Fig. 1C), which were most noticeable at E10.5, when the rate of neuronal differentiation is highest in the pMN (Kicheva et al. (2014)). Nevertheless, the average area of cells assayed in this way was consistent with previous measurements (Kicheva et al. (2014)). Using these data we set out to develop an in *silico* model of the neuroepithelium.

### A vertex model of the neuroepithelium including inter-kinetic nuclear movement

To investigate the influence of mechanics on neural progenitor geometry we constructed a 2D vertex model of the apical surface of the neural tube. In these simulations, cells are represented as polygons. The behaviour of each cell is governed by the movement of its vertices that follow a deterministic overdamped motion given the energetic contributions of cell elasticity, junctional forces arising from cortical contractility of a cell and the effect of cell-cell adhesion and cortical tension (Landsberg et al. (2009); Farhadifar et al. (2007); Aegerter-Wilmsen et al. (2010); Fletcher et al. (2014); Honda and Nagai (2015); Alt et al. (2017)). For the purposes of the simulation we developed custom Python code using an Euler method to solve the movement equation of each vertex, Eqn (3).

Topologically, the neural tube is a cylinder that grows at different rates along its anteroposterior and dorsoventral axes (Fig. 2A). Our analysis focuses on a region along the AP axis at the forelimb level. To take this into account, we introduced periodic boundary conditions in the AP axis by simulating the neural tube as a torus. For visualisation we unwrapped the torus by cutting along both DV and AP axes to allow simulations to be rendered in 2D (Fig. 2A, bottom).

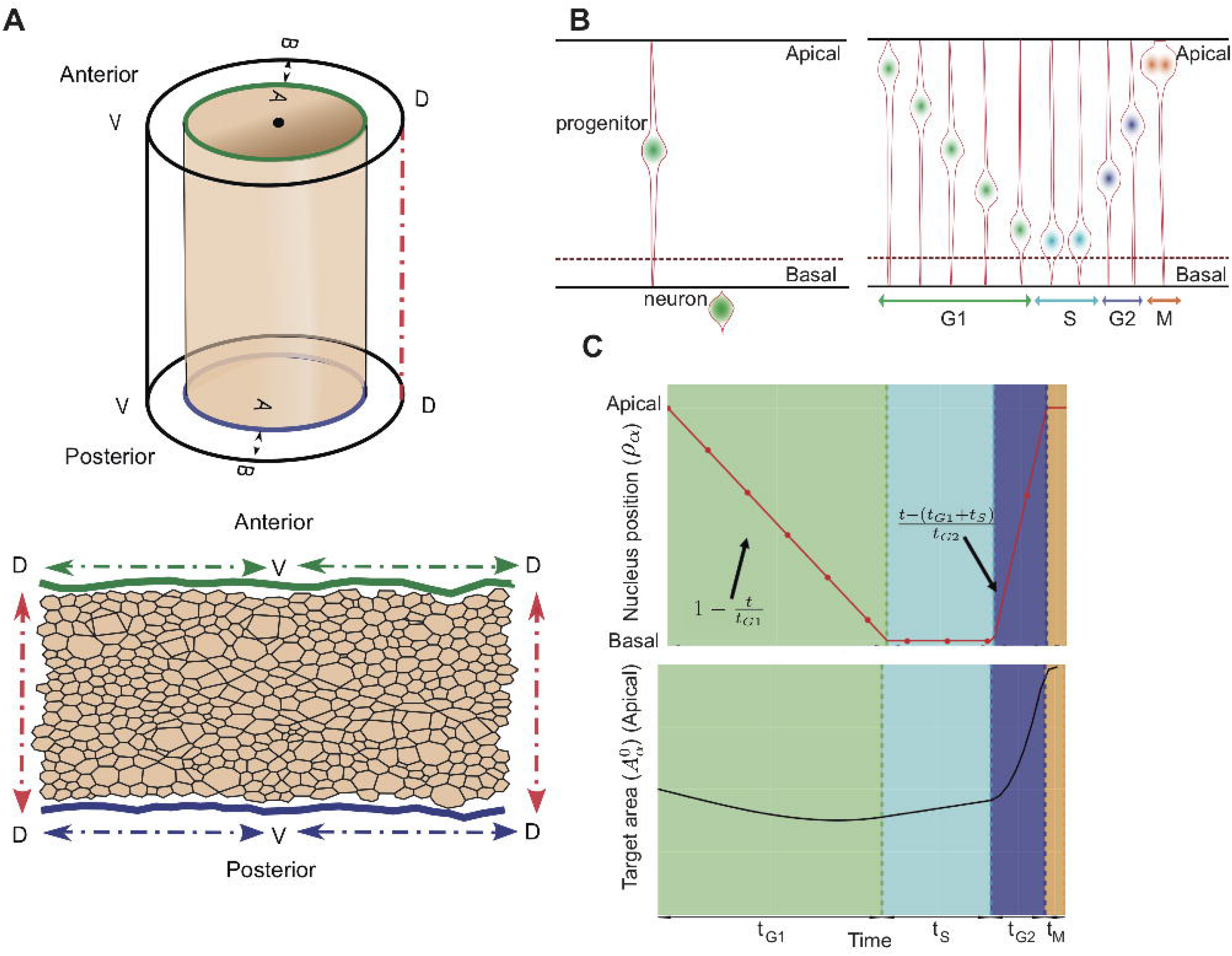

To describe the behaviour of neural progenitors within the simulation, a detailed description of the cell cycle is required including cell growth, division and differentiation. Upon neuronal differentiation, the cells delaminate from the epithelium by losing their apical attachments and extruding basally (Fig. 2B, left). During each cell cycle, progenitors in the neuroepithelium undergo IKNM. This involves the translocation of the nuclei and the bulk of the cell volume along the apical-basal axis of the neuroepithelium. Mitosis occurs at the apical side of the epithelium. Nuclei move basally as they proceed through G1 and undergo S-phase towards the base of the epithelium. During the G2-phase, nuclei migrate back to the apical surface for mitosis. A consequence of IKNM is that the apical area of cells, corresponding to the surface represented in the simulations, is affected by the cell cycle stage. When a cell enters mitosis, the nucleus translocates towards the apical surface and the cell rounds up. Thus, cells that approach mitosis expand their apical area and compress neighbouring cells. As a consequence, cells are likely to achieve their largest apical surface area in late G2 and M phase, and their smallest surface area in S-phase. The measured duration of cell cycle phases (Kicheva et al. (2014)) (Table 1), can therefore be used to derive an approximation for the temporal changes in apical surface area of cells caused by IKNM.

**Table 1.**
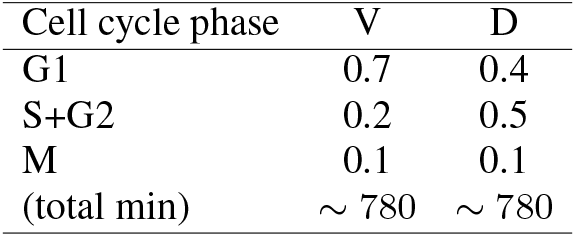
Proportion of cells in the indicated cell cycle phases and total cell cycle time in minutes at stage E10 of cells in the ventral (V) and dorsal (D) region of the neural tube. Data from (Kicheva et al. (2014)).

| Cell cycle phase | V | D |
| --- | --- | --- |
| G1 | 0.7 | 0.4 |
| S+G2 | 0.2 | 0.5 |
| M | 0.1 | 0.1 |
| (total min) | $\sim 780$ | $\sim 780$ |

To accommodate the effect of IKNM in our simulations we introduced a time-dependent target area function, 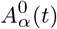 in Eqn (4), which describes the desired apical area of cell. This function depends on the age of the cell and the cell-cycle phase and was constructed to account for the measured cell cycle dynamics:

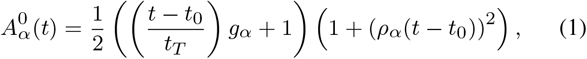

where *g_α_* is the growth rate of the cell *α, t*_0_ is the moment when the cell *α* is born, *t_T_* is the total time of the cell cycle and *ρ_α_* (*t* − *t*_0_) represents the apical-basal position and depends on the phase of the cell-cycle by a piece-wise linear function incorporating the dynamics of the cell-cycle:

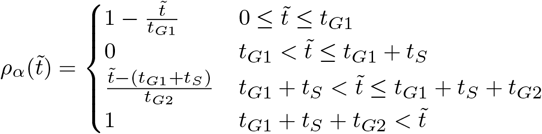

where *t*_*G*1_, *t_S_* and *t*_*G*2_ are the respective cell cycle phase durations and 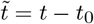. The position of a cell body along the apicobasal axis (see scheme in Fig. 2B, right) is given by the function *ρ_α_* (*t*) where apical is 1 and basal is 0. The function *ρ_α_* (*t*) is defined by four different straight lines which correspond to each cell-cycle phase (Fig. 2C). In G1, the nuclear movement is from apical to basal and takes *t*_*G*1_ time and thus decreases linearly with time at rate 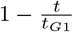. During S-phase, the nucleus stays basal for time *t_S_*, and *ρ_α_* (*t*) is set to 0. Basal to apical migration occurs during G2, over the period *t*_*G*2_, and is represented by the increasing function 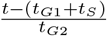. During mitosis, the function takes value 1. The functional form of *A_α_* (*t*) is the product of a term which grows linearly in time, as we assume the volume of the cell does, and a term which interpolates between 1/2 and 1, as the fictive cell moves from the basal to the apical surface, with a higher rate of increase as it approaches the latter surface.

Implemented in this way, the target area of a cell, which describes the desired apical surface area of cell, takes account of both the growth of a cell during the cell cycle and the position of the cell body along the apicobasal axis. It results in the apical area of a cell slowly reducing during G1, corresponding to the cell body moving from the apical to the basal surface at the same time as the cell is growing, then beginning to increase slowly during S phase, rapidly expanding during G2, as the cell returns to the apical surface for division and growing slowly during mitosis (Fig. 2C, bottom).

Cell division occurs when the cell cycle has been completed and the volume of the cell exceeds a critical value, *A_c_*. As a cell undergoes division, two new vertices are created to form a new edge, the location of one end of this new edge is chosen as the midpoint of a randomly selected edge of the dividing cell with probability proportional to the edge length. The other end is the midpoint of the opposite edge, if the cell has an odd number of sides the second edge is the closer mid edge. The newly generated sister cells then commence the next cell cycle.

In the neural tube, newly generated neurons lose their apical attachments, delaminating and migrating basally (Fig. 2B, left). Hence, neuronal differentiation leads to the loss of cells from the plane of the neuroepithelium. In the simulation, this is achieved by identifying cells committing to differentiation, suppressing growth in these cells by assigning their target area equal to zero and allowing their area to decrease. As a cell’s area drops some of its edges become small and disappear under certain T1 transition conditions (see below), this ultimately results in elimination of the cell. At the stage of development we are modelling, cells differentiate predominantly within the pMN domain. In simulations, we select cells to differentiate with a fixed probability per unit time.

The combined effect of cell growth, division and differentiation results in cells moving relative to each other, producing local remodelling of the epithelium and rearrangements of neighbouring cells. In the simulations these topological rearrangements occur through T1 transitions. During a T1 transition an edge with a length shorter than T1 (chosen to be 3% of the average edge length in the tissue) is eliminated and a new edge of length *l_new_* expands perpendicular to the old edge (values given in Table 2). However, if the rearrangement results in the formation of a two-sided cell, the cell is removed from the epithelium.

**Table 2.** Simulation constant parameters.

| Parameter | Meaning | Value | Ref |
| --- | --- | --- | --- |
| $K$ | Elasticity coefficient | 1 a.u. | (Farhadifar et al. (2007)) |
| $A_c$ | Critical area | $30 \mu m^2$ | - |
| $T_1$ | Length threshold | $0.048 \mu m$ | (Kursawe et al. (2018)) |
| $l_{new}$ | Distance new edge nodes after $T_1$ | $1.01 T_1$ | (Kursawe et al. (2018)) |
| $\Delta t$ | Time step | $0.46 s$ | (Fletcher et al. (2013)) |
| $\mu$ | Medium viscosity | $0.276 K \mu m^2 s$ | - |
| $\mu'$ | DV drag viscosity | $0.276 K \mu m^2 s$ | - |
| $\mu''$ | AP drag viscosity | $13.8 K \mu m^2 s$ | - |
| $\lambda$ | Proliferation rate | $0.05 h^{-1}$ | (Kicheva et al. (2014)) |

### Simulation parameter estimation

We next used the experimental data to identify model parameters for which simulations match *in vivo* observations. The dynamics of the simulations are determined by the Hamiltonian (Eqn (4)) that takes into account the energetic contributions of different cellular mechanical properties. The minima of this Hamiltonian can be described using two dimensionless parameters: 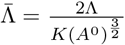 and 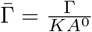 (Farhadifar et al. (2007)). In the standard implementation of the model this leads to a phase diagram describing four different parameter regions where the tissue has different biophysical properties, (SM II and (Magno et al. (2015); Farhadifar et al. (2007))). The Hamiltonian we use differs from standard implementations in that the target area term includes a cell-cycle dependent component. However, since vertex movement is substantially faster than the cell cycle, the same phase diagram remains applicable (for more details of the derivation of the phase diagram see SM II).

We focus our attention on the region of the phase diagram exhibiting epithelial properties (Regions II and III in Fig. SM2); this is given by the following relation between normalised tension and contractility parameters, (Magno et al. (2015)):

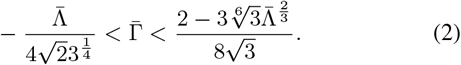

To narrow down the region of parameter space, 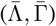, relevant for neural tube simulations, we systematically screened parameter sets to identify those that generated cell geometries comparable to experimental data. We compared experimental and simulated empirical cumulative distribution functions (ECDF) of area, perimeter, standard deviation of area and perimeter, number of cell sides, and cell elongation. We used experimental data from E11.5 dorsal neural tubes compared against the ECDF obtained from vertex model simulations with different combinations of 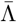 and 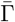.

Assessing the match between experimental and simulation data indicated a diagonal region in parameter space for which all the measured features of the *in silico* cell geometries closely matched those observed *in vivo* (Fig. 3A). The shape of this region is similar to previously published vertex model simulations (Kursawe et al. (2015)). Moreover, the agreement with experimental data was better in the model with IKNM (Fig. 3A), compared to a standard model formulation without IKNM in which the target area is constant over time (Fig. S.1). Therefore, in subsequent simulations we used the model with IKNM and 6 parameter sets selected from different locations from within the region of parameter space representing the best agreement with experimental data.

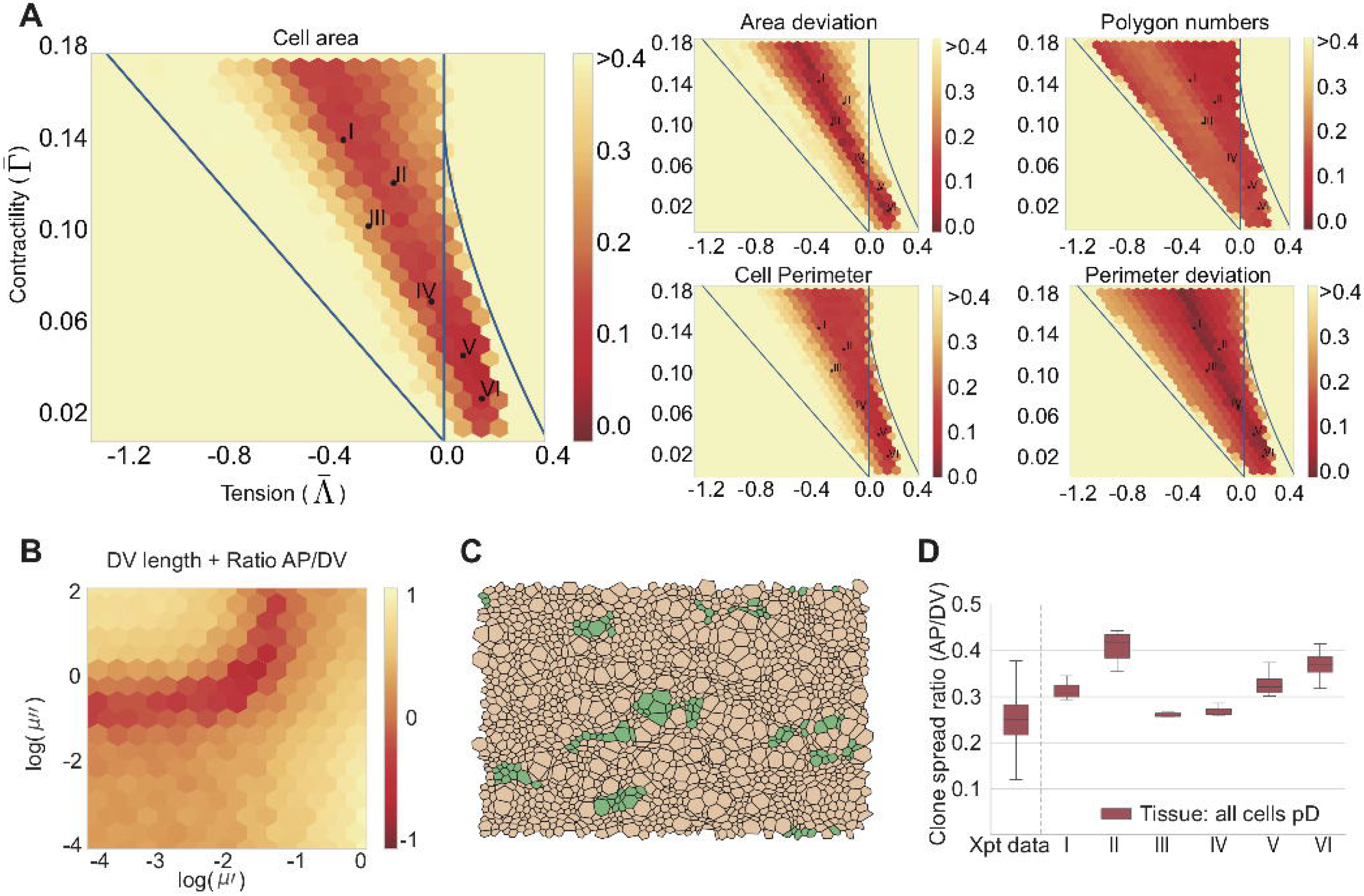

### Simulating anisotropic tissue growth

We next turned our attention to the overall growth of the tissue. Our previous experimental studies (Kicheva et al. (2014)) indicated that the tissue grows asymmetrically in DV and AP directions. During the period under consideration, the dorsoventral length of the tissue increased ~3 fold more than the anteroposterior length (Kicheva et al. (2014)). This effect on the tissue aspect ratio (AP/DV) was reflected in the shape of lineage-related clones of cells, such that the mean ratio of AP to DV spread of the clones outside of pMN domain was ~ 0.3.

In the simulations, expansion along dorsoventral and anteroposterior axes is resisted by drag forces that have coefficients, *μ*′ and *μ*″, respectively. A difference between these two coefficients generates different rates of dorsoventral and anteroposterior growth during development (see SM I for more information), imitating the effect of physical constraints on *in vivo* tissue expansion. For all of the 6 selected parameter sets, we found that experimentally observed aspect ratios were recovered for values *μ*’ ~0.02 and *μ*″ ~1 (Fig. 3B and Fig. S.2A).

To test the effect of these asymmetric forces, we examined the shape of clones in simulations by tracking lineage-related cells *in silico*. For this, simulations were started from a field of 100 cells and run for 30 hours of biological time to allow the simulation to equilibrate. Following this initialisation period, the progeny of individual cells were tracked for a further simulated 48h. This corresponds to an average of 3-4 cell divisions, mimicking the experimental conditions in which the *in vivo* clonal data were generated. Similar to the experimental data, in *silico* lineage-related cells (clones) tended to form coherent groups and the shape of clones was similar between experiments and simulations (Fig. 3D). For all 6 parameter sets, cells within a clone tended to spread more along the DV axis compared to the AP axis to give an *in silico* AP to DV aspect ratio of ~ 0.3 (Fig. 3D), similar to clones in the mouse neural tube (Kicheva et al. (2014)). Taken together this suggests a good correspondence between the behaviour of cells in the simulation and those in the real neuroepithelium.

### The rate of neuronal differentiation affects the shape of progenitor clones

Having established a simulation framework and identified parameters that mimicked neuroepithelial behaviour we set out to address whether differences in differentiation rate could affect the spatial allocation of cells and shape of clones. Whereas in the dorsal progenitor domains (pD), the spread of clones is elongated along the DV axis compared to the AP axis, within the motor neuron progenitor domain (pMN), clones had an average AP/DV aspect ratio of ~ 1 indicating equal spread in DV and AP directions. As a consequence, this domain grows isotropically. Progenitors within the pMN differentiate at a substantially higher rate than other progenitors at this stage of development (Kicheva et al. (2014)) raising the possibility that this accounts for the difference in clone shape.

We implemented a pMN domain in simulations by defining a region of tissue with an appropriate differentiation rate. Following the initialisation period, a pMN domain comprising 30% of DV length of the neural tube was introduced into the simulations by imposing a differentiation rate of 0.1*h*^−1^ on cells in this region, corresponding to the differentiation rate of motor neurons *in vivo*, (Kicheva et al. (2014)). The remainder of the tissue was designated the pD domain and lacked differentiation, representing the much more slowly differentiating dorsal progenitor domains *in vivo*. Simulations were continued for a further period equivalent to 48h of biological time, see Fig. 4A.

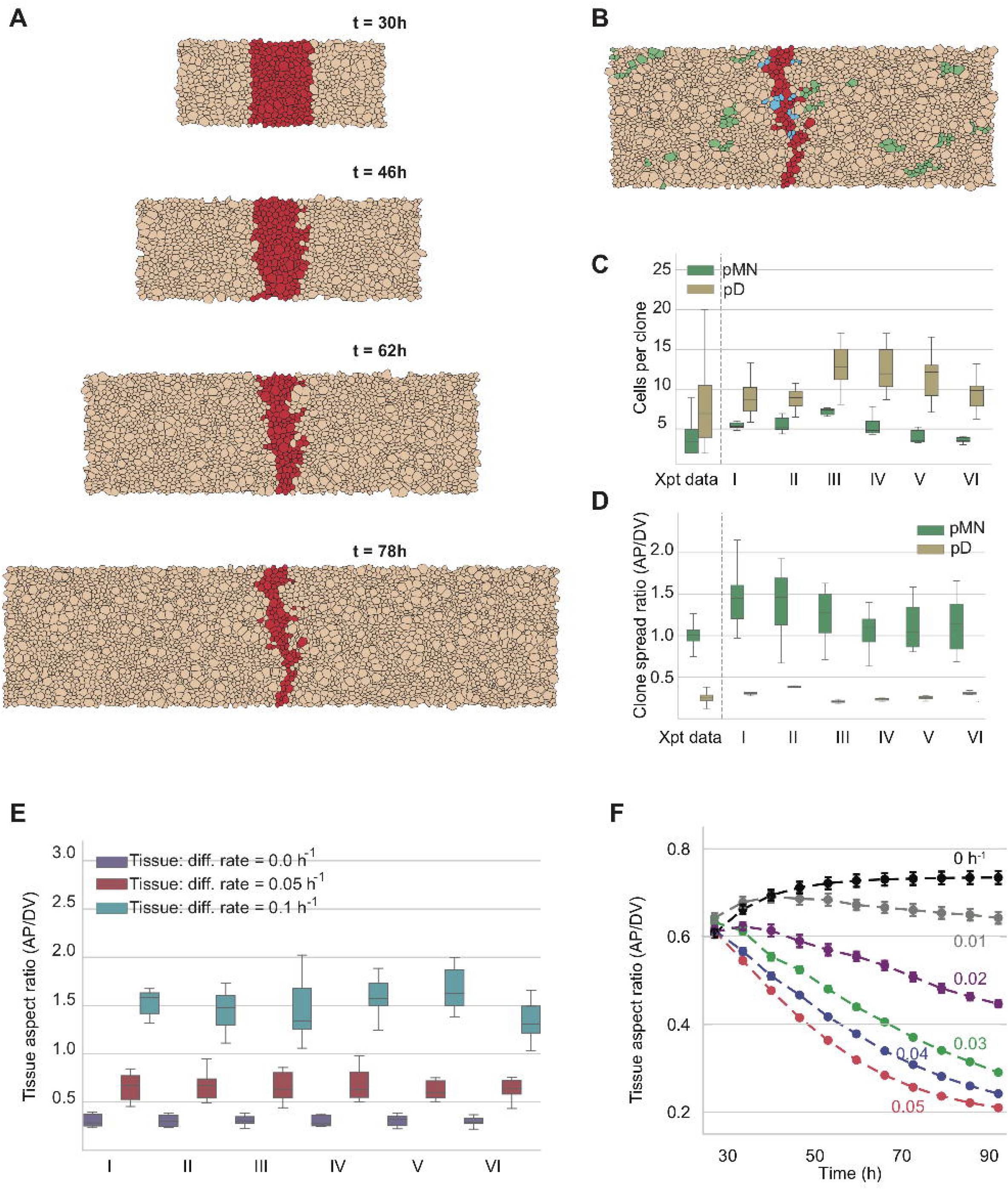

At the end of the simulations, the size of the pMN and pD domains and number of cells per clone in the two domains were measured. In all 6 mechanical parameter sets, the proportion of tissue comprising the pMN decreased from the initial 30% DV length of the tissue to ~ 5% DV length (Fig. S.2A). This is a consequence of the increased differentiation rate resulting in a loss of progenitors from the pMN. This decrease in the DV extent of the pMN matches the experimentally observed reduction in the DV proportion of the neuroepithelium occupied by the pMN domain from 30% of the neural tube at E9 to 5% 48h later at E11 (Kicheva et al. (2014)). Moreover, clones in the pD domain were comprised of 8-12 cells on average, consistent with an average of 3-4 cell divisions that occur in the 48h period (see Fig. 4B and Supplementary Movie1). By contrast, pMN clones contained 4-5 cells per clone. These *in silico* clone sizes are consistent with the clone sizes observed in the experimental data (Fig. 4C). Together these data indicate that the behaviour of the simulated pMN and pD domains matches the behaviour observed *in vivo*.

We then examined the spread of clones along the AP and DV axes (Fig. 4D). Similar to the simulations lacking a pMN domain (Fig. 4D), clones within the pD region were anisotropic with an AP/DV aspect ratio of ~ 0.3. This matches experimental data (Fig. 1A, (Kicheva et al. (2014))). By contrast, for all 6 parameter sets, clones within the pMN domain had a substantially higher AP/DV aspect ratio (Fig. 4D), consistent with the experimentally measured aspect ratio of pMN clones of 1 ± 0.3 (Kicheva et al. (2014)). These results reveal that the difference in the shape of clones in the pMN compared to the rest of the neural tube can be explained by the increased differentiation rate of these cells.

### The anisotropy of the tissue expansion rate growth depends on tissue expansion rate

To investigate how the increased rate of differentiation affects the anisotropy of growth, we analysed how the tissue anisotropy changed when different differentiation rates (0-0.1 *h*^−1^) were imposed uniformly throughout the tissue. This revealed a relationship between differentiation rate and the aspect ratio of the tissue: the higher the differentiation rate, the larger the AP/DV aspect ratio of the tissue (Fig. 4E). This effect is consistent with the observation that higher differentiation rate correlates with higher AP/DV aspect ratio of clonal shape (Fig. 4D, Fig. S.2).

The decreased anisotropy observed in the pMN domain might result from the decreased net growth rate of the pMN, rather than directly from the increased differentiation. To investigate the effect of tissue growth rate on anisotropy, we began by further simplifying the problem and constructing a simple model in which we assumed that the internal boundaries of the tissue could rearrange instantaneously (see SM III). This predicted that the aspect ratio tends asymptotically to a value that depended on the drag coefficients and the growth rate. For very slow growth, the aspect ratio would be close to one, whilst for very rapid growth, it would be close to the square root of the ratio of drag coefficients. Thus the effect of the drag on tissue anisotropy would be less pronounced for slow growth rates, leading to more isotropic growth. To test this hypothesis, we ran simulations without differentiation but with varying proliferation rates (Fig. 4F). Consistent with our hypothesis, the slower proliferation rates decreased the anisotropy of tissue growth. Thus, increased differentiation *per se* was not necessary for the observed behaviour. Instead, the net growth of tissue affects its aspect ratio.

Further investigation showed that the effect of differentiation rate on the aspect ratio of the tissue was more complex than simply slowing the effective tissue growth. Whilst both increasing the differentiation rate and decreasing the proliferation rate correlated with decreasing anisotropy at a given time (see Fig. S.2A,F), the anisotropy that corresponded to a specific net growth rate was not always the same. Instead, different degrees of anisotropy were achieved for the same net growth rate, depending on whether proliferation or differentiation was modulated (Fig. S.2F). Furthermore, tissues of similar size (generated by similar net growth rates) would adopt different aspect ratios depending on the relative contributions of proliferation vs differentiation to the net tissue growth (Fig. S.2A-F). Tissues that differentiate have increased anisotropy (lower AP/DV ratio) compared to tissues that reach the same size without differentiation (compare Fig. S.2G and H). We postulate that increasing differentiation rate facilitates the rearrangement of internal boundaries which allows the tissue to tolerate more anisotropy. Thus, differentiation has opposing effects: i) it slows growth, which tends to make growth more isotropic and ii) it facilitates internal boundary rearrangements, which tends to allow more anisotropy. Further work will be needed to fully understand the determinants of tissue aspect ratio.

We next turned our attention to the cellular dynamics that result in the anisotropic growth of the tissue. We first measured the orientation of T1. In the pD domain, these more frequently resulted in topological rearrangements that replace an AP directed edge with one in the DV direction (Fig. 5A). Such transitions cause cells to intercalate, expanding the DV axis. There was no such bias in cells of the pMN domain. A consequence of these dynamics was a change in the overall shape of cells. While the cells were equally eccentric in the pMN and pD domains, in the pD domain cells tended to be more elongated along the DV axis, whereas cells in the pMN domain tended to be orientated with their long axis in AP direction (Fig. 5C). Strikingly, these changes in cell orientation were also observed in the experimental data, where pD cells were more elongated in DV direction, whereas cells from the ventral half of the neural tube were more elongated in the AP direction (Fig. 5B, C).

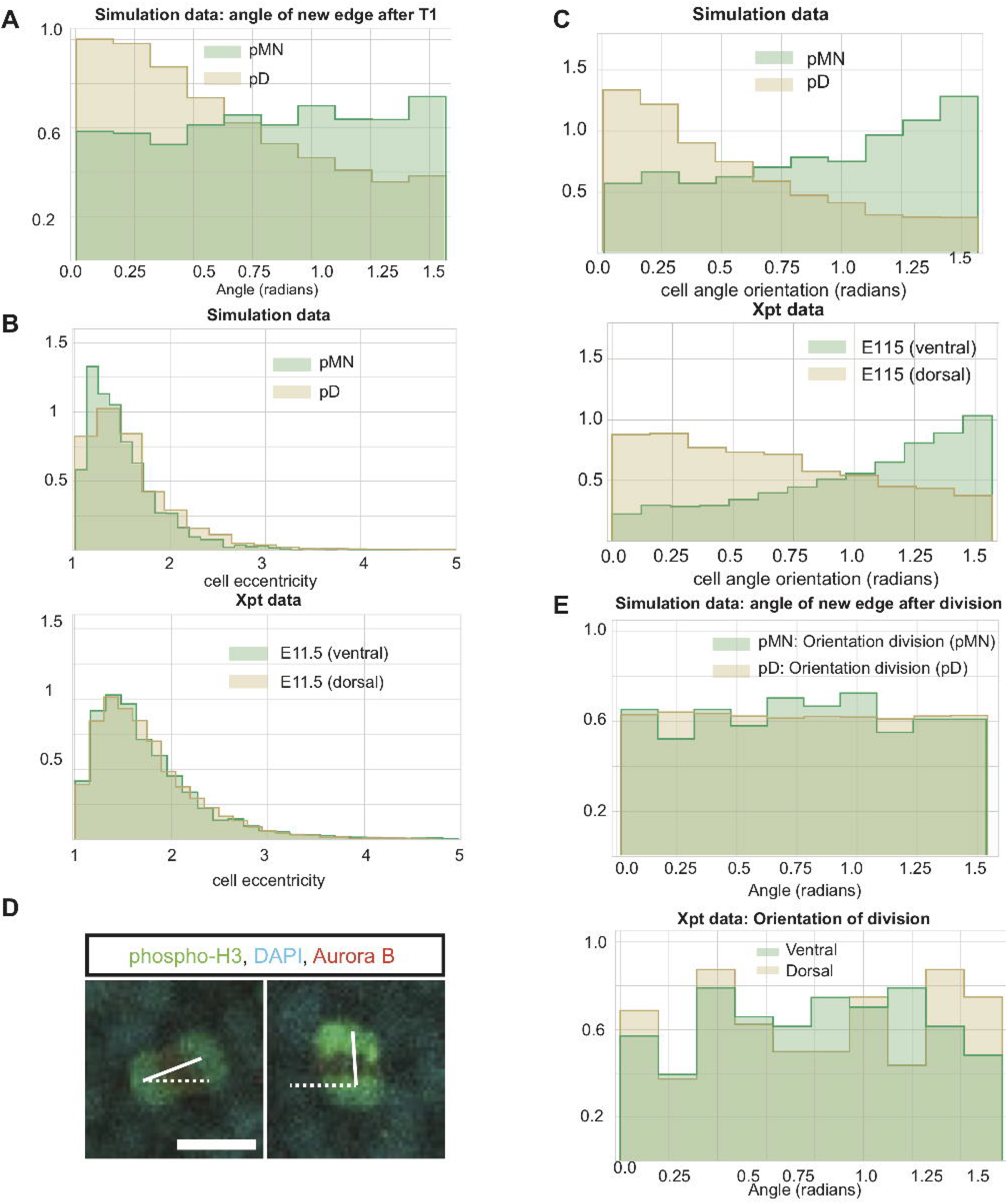

In contrast to the bias in cell rearrangements, the axes of cell divisions in our simulations were distributed uniformly in both the pD and pMN domains (Fig. 5D, E) indicating little, if any, bias in division orientation. In the simulations this is a consequence of mitotic cells markedly reducing their eccentricity, allowing for random orientation of the division angle (Fig. S.2). To test whether this was consistent with the *in vivo* observations we examined the orientation of mitotic spindles in anaphase cells, as a proxy for the orientation of cell division at E10.5 (Methods). This revealed a uniform distribution of cell division orientation in both dorsal and ventral regions of the neural tube (Fig. 5E). Together these results suggest that a difference in cell rearrangements, rather than oriented cell division, account for the reduction of anisotropic growth in the pMN domain.

In summary, the experimental observations are consistent with a model in which tissue growth is resisted by forces which are larger in the AP direction, causing anisotropic growth. This effect is lessened in slow-growing epithelia. Thus, clones in the rapidly growing pD domain become more anisotropic after a fixed period of time than clones in the more slowly growing pMN domain. When the tissue grows anisotropically, it does so by biasing the direction of T1 transitions, rather than biasing the orientation of cell divisions.

## DISCUSSION

To understand the mechanisms by which tissue pattern, mechanics and growth are coupled in the vertebrate neural tube we used experimental data to construct a mechanical model of the developing neuroepithelium. This allowed us to define and test how proliferation and differentiation of individual cells, together with global mechanical constraints, influence the spatiotemporal dynamics of pattern formation in the tissue. Previous observations indicated that there are differences in the anisotropy of clonal shape within the plane of the epithelium in different dorsoventral regions of the neural tube (Kicheva et al. (2014)), however, how this anisotropy emerged was not understood. Our simulations and analysis indicate that the differences arise as a consequence of mechanical constraints on tissue growth acting with differences in the local rate of neuronal differentiation, leading to different rates of cell loss from the epithelium that affects the net growth of the tissue. Comparisons with experimental data were consistent with this interpretation. The analysis suggested an explanation for how an isotropic process, such as cell differentiation, can produce an anisotropic outcome, the direction of clonal expansion.

We adopted the well-established vertex model framework to describe neuroepithelium growth (Nagai and Honda (2001); Fletcher et al. (2013); Asgari-Targhi (2012); Smith et al. (2011); Chiou et al. (2012); Canela-Xandri et al. (2011); Bock et al. (2010); Farhadifar et al. (2007)). This approach provides a scalable and computationally efficient means to understand how tissue morphogenesis is influenced by the combined effect of cell shape, forces generated by growing cells and external mechanical constraints. However, one of the challenges of modelling the neural tube epithelium resides in the 3D dynamics of neural progenitors. Similar to many pseudostratified epithelia, cells within the neuroepithelium undergo interkinetic nuclear movement (Sauer and Walker (1959)) in which cell nuclei migrate between the apical and basal surfaces in synchrony with the cell cycle. Whilst previous approaches focussed on a purely 2D representation (Farhadifar et al. (2007); Aegerter-Wilmsen et al. (2010)), here we extended the formalism by including the effect of the IKNM and cell cycle on the preferred target apical area. This allowed the fitting of mechanical parameters from experimental images of the apical plane, without requiring 3D reconstruction of the neural tube. We could thus recapitulate more faithfully the pseudostratified dynamics of the tissue without compromising the computational efficiency of the model.

To identify the mechanical parameters of the model, we varied the tissue tension, contractility and dissipative forces and compared descriptors of cell and tissue geometry in the resulting simulations to experimental data. The parameters we identified are in line with those used in previous epithelial vertex models, such as the wing disc (Farhadifar et al. (2007)). The parameter values that produced the highest correlations with experimental data reside in the region of parameter space in which the unperturbed ground state is represented by hexagonal packing (Nagai and Honda (2001); Farhadifar et al. (2007); Magno et al. (2015); Gibson et al. (2006)) and there is a negative correlation between tension and contractility. This is as expected (Kursawe et al. (2015)) and is consistent with an overall similarity of cell shapes and behaviours in different epithelia. Nevertheless, quantitative details, such as cell areas and geometries, differ and are accounted for by the values of the parameters used. Moreover, a model without IKNM yielded a different set of parameters (Fig. S.1A) that fit the data less well. Although the distribution of cell orientations was comparable to experimental data for both models (Fig. S.1B, left), the simulations with IKNM reproduced the experimentally observed cell shapes more accurately, probably because mitotic cells are rounder as a result of IKNM (Fig. S.1B, right).

Our previous analysis (Kicheva et al. (2014)) of clone shape *in vivo* revealed that, in most of the tissue, the neural tube grows faster in DV than in AP direction. We found that cell divisions do not show a preferred orientation in the plane of the epithelium, hence this anisotropy of tissue growth must arise from mechanical constraints. To model this, we assumed that the overall growth of the tissue was resisted by drag forces with different coefficients in the two directions. Screening systematically these coefficients, we identified the values that produced clone shapes similar to experimental data.

The sources of the resistive forces in epithelium are generally poorly understood (Alt et al. (2017)) and in the neural tube their basis remains to be determined. It is possible that neural tube expansion is mechanically constrained by the adjacent tissues. Thus, the laterally located somites are likely to affect radial expansion, whereas the process of axis elongation, which results from addition of cells to the caudal end of the neural tube and cell motility in the posterior of the embryo (Bénazéraf et al. (2010); Mongera et al. (2018)), might contribute to the forces acting in the AP direction. Obtaining mechanical measurements of the mouse neural tube in *situ* might provide insight, however accessing and assaying these properties *in vivo* without surgical disruption is difficult or impossible. Methods for the *ex vivo* culture of embryos, or dissected embryonic tissue, in which specific mechanical constraints can be applied and measured, might offer an alternative investigative approach.

A limitation of our approach is the lack of kinematic data on cell shape and movement. Instead we relied on static images and experimentally inferred cell dynamics. The inaccessibility to live imaging of the unperturbed apical surface, which forms the internal face of the intact neural tube, remains a challenge in this regard. While it is conceivable to develop imaging preparations that would allow apical imaging of neural progenitors, these would most likely involve mechanically disrupting the neural tube with the risk that epithelial morphogenesis is affected.

Encouraged by the similarity between simulations and experimental data we used the model to examine the clonal spread in different progenitor domains in the neural tube. Although the shape of clones was anisotropic throughout most of the neural tube, clones in the pMN domain were smaller and rounder (Kicheva et al. (2014)). pMN cells are distinguished by their high rate of terminal differentiation (Sagner et al. (2018); Kicheva et al. (2014); Ericson et al. (1992)), which causes the loss of cells from the neuroepithelium and consequently smaller clone sizes in the pMN domain compared to other domains. However, the basis for the difference in clone shape was unclear. pMN cells are molecularly distinct from other progenitors and one possibility was that cell orientation or arrangement was under molecular control. For instance, pMN progenitors express different sets of adhesion molecules than their adjacent domains (Rousso et al. (2012)), raising the possibility that cell-cell communication plays a role in shaping the pMN domain.

Strikingly, the model showed that the anisotropy observed experimentally could be reproduced without invoking mechanisms that directly control the mechanical properties of pMN progenitors or division orientation. Instead, differences in clone shape emerged as a consequence of the increased differentiation rate of pMN progenitors in conjunction with the global difference in the resistive forces in AP and DV directions. The difference in resistance causes the tissue to become increasingly more anisotropic with time. Futhermore, increasing the differentiation rate or decreasing the proliferation rate causes the tissue to grow more slowly and become less anisotropic over a given period of time. Hence, the net growth rate is a governing factor that influences the degree of anisotropy, with slow growth being more isotropic. We found that this change in anisotropy over time not only depends on the overall growth rate of the tissue, but also on the relative magnitudes of the proliferation and differentiation rates. Thus, for any growth regime (with fixed rates of proliferation and differentiation) there is a characteristic temporal trajectory of the anisotropy of growth. In particular, increased differentiation, which removes cells from the epithelium, effectively increases the fluidisation of the tissue (Ranft et al. (2010)). This facilitates cell rearrangements and alters the degree of anisotropy that is achieved for a given tissue size compared to growth without differentiation. In summary, the difference in the growth regime between domains influences the degree of anisotropy. This highlights how changes in isotropic processes, such as differentiation and proliferation, can result in a change in the shape of clones and affect the isotropy of tissue growth. We note that detailed information about the forms of the resistive forces in the two directions is not available and this warrants further investigation.

Our data indicate that the shape of clones become progressively more elongated with time, mimicking the degree of tissue anisotropy. Consistent with this, in regions of low differentiation, T1 transitions, which occur when the length of an edge falls below a threshold, were preferentially oriented such that it was more likely for an edge in the AP direction to be replaced by one in the DV direction (Fig. 5A). The cell intercalation that results fromT1 transitions contributes to the tissue extension in the DV direction. This is similar to the way in which tissue domains elongate through convergent extension, for example during gastrulation in the frog (Keller et al. (2000)), although in the case of the neural tube instead of solely rearranging the cells, their number also increases by proliferation and consequently the tissue expands along both AP and DV axes.

Our analysis shows that the orientation of cell divisions in experimentally observed tissues, as well as in simulations, is random. This suggests that cell division orientation does not contribute to anisotropic tissue growth (Li et al. (2014)). A surprising observation in our data is that despite preferential cell elongation in DV direction, cell division orientation is apparently random in the AP/DV plane, this could be explained by the decrease in eccentricity observed in mitotic cells as a result of IKNM (Fig. S.2). Cell division has been found to be frequently oriented along the longest planar axis of a cell (Mao et al. (2011); Baena-López et al. (2005); Bosveld et al. (2016); Seldin and Macara (2017)). While in some systems the orientation of cell division is determined by external or internal cues independent of cell shape. Prominent examples include the planar cell polarity pathway (Gong et al. (2004)), or determinants of asymmetric cell division, such as NuMa, Par3, LGN, Pins etc. (Bowman et al. (2006); Konno et al. (2007); Gillies and Cabernard (2011)). In neural progenitors in the spinal cord, the plane of cell division is regulated along the apicobasal axis and is accompanied by extensive rotations of the metaphase plate (Morin et al. (2007)). This regulation is important for maintaining the integrity of the epithelium. Our data are consistent with the idea that the apicobasal orientation of the spindle is the dominant mode of regulation in the spinal cord, with no specific mechanism acting to orient the planar angle. It could be that the random orientation of divisions in the epithelial plane and the decoupling from cell shape is necessary to achieve efficient apicobasal orientation (Morin et al. (2007)). Further studies will be necessary to investigate this thoroughly.

The increased rate of differentiation of pMN progenitors, which drives the distinct morphodynamics of the pMN, depends on the transcription factor Olig2 within this domain. By repressing Notch target genes, Olig2 promotes neurogenesis (Sagner et al. (2018)). A consequence of the difference in clone shape between pMNs and other progenitor subtypes is that cells in all progenitor domains expand at equal rates along the AP axis, despite the overall smaller size of pMN clones. Thus, there is no net AP movement between progenitor domains and cells stay in register as development proceeds. This means that cells in different DV domains with the same AP identity remain adjoining. Since AP identity is established early during neural development, maintaining position relative to other cells in the epithelium may be important for the later assembly of functional neuronal circuits. The guidance of axons, identification of correct partners, and subsequent synaptogenesis depends on local cues within the neuroepithelium, hence precision depends on cells initially residing in the appropriate location. As well as influencing the rate of MN formation, Olig2 is also a component of the gene regulatory network responsible for the initial DV positioning of the pMN within the neural tube (Novitch et al. (2001)). Together, therefore, Olig2 plays a pivotal role in the spatial and temporal organisation of motor neuron specification and differentiation.

In conclusion, we have described a vertex model of a pseudostratified epithelium and used it to study the growth of the neural tube and how cell differentiation influences clone shape. In future work, we wish to couple this tissue model to quantitative descriptions of the spread of morphogens that pattern the tissue and the gene regulatory networks that specify neuronal subtype identity. In this way, we hope to gain insight into the coupling of growth and patterning in the neural tube and understand how the position, precision and proportions of cell types is achieved.

## MATERIALS AND METHODS

### Experimental data analysis

E10.5 and E11.5 mouse embryos were collected and processed for dissection, fixation, immunostaining and flat-mounting as previously described (Kicheva et al. (2014)). Primary antibodies used were: mouse anti-ZO-1 (33-9100, Zymed labs), rabbit anti-phospho Histone H3 (Novus Biologicals), mouse anti-Aurora B (AIM1, BD Transduction), rabbit anti-Olig2 (Millipore), mouse anti-Pax3 (DSHB). For the analysis of clonal shape, we reanalysed the data in (Kicheva et al. (2014)), in which YFP expression was sparsely induced within the neuroepithelium at E9.5 and assessed at E11.5 of development. The AP and DV spread of the clone were defined as the x (AP) and y (DV) components of first eigenvector of the second moment matrix of the coordinates of cells comprising the clone (for details, see (Kicheva et al. (2014))).

To measure the orientation of cell division, E10.5 embryos were stained with DAPI, phospho-H3, and Aurora B to mark dividing cells, and Pax3 and Olig2 to mark the dorsal and pMN domains, respectively. Only cells in anaphase, which have low levels of phospho-H3, separated sister chromatids and Aurora B staining associated with the central spindle, were considered for the analysis (Fig. 5B, bottom). The orientation of the chromosomes with respect to the anteroposterior axis of the embryo was measured.

For the analysis of cell geometries, we used images of flat mounted embryos of approximate size 80 × 80 *μ*m. Images taken within the dorsal half are comprised of Pax7+ pD progenitors, while images taken in the ventral part of the neural tube can contain up to 50% pMN cells, and the rest intermediate progenitor identities (p2-p0). Images were processed using the Fiji plug-in ‘Packing Analyzer v2.0’ (Aigouy et al. (2010)), which segments the image, classifies cell edges and vertices, and measures cell areas, perimeter, neighbours. Segmentation mistakes were manually corrected.

### Vertex model description

Cells are represented as polygons with straight edges connecting vertices. Cells are enumerated by *α* = 1,…,*N_c_* and vertices are enumerated by *i* = 1,…,*N_v_*. The evolution of each cell in these models is governed by the motion of its vertices, which are typically assumed to obey deterministic equations of motion. It is usual to make the simplifying assumption that the motion of vertices are over-damped (Drasdo (2000)), and inertial terms are small compared to dissipative terms. This leads to first-order dynamics. The evolution of the position *r_i_* of vertex *i* determined by:

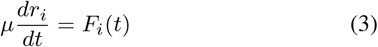

where *F_i_*(*t*) denotes the total force (except drag) acting on vertex *i* at time *t* and *μ* denotes its drag coefficient. The main difference between models lies in the definition of the force *F_i_* that can be derived from an energy function, *E_i_*, which includes the different cell-cell interactions. In our model we will use a modification of the energy function described in (Farhadifar et al. (2007))

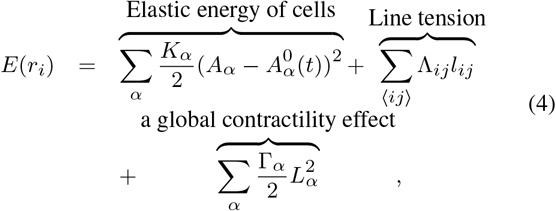

for which 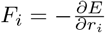.

The first term describes an area elasticity with elasticity coefficients *K_α_*, for which *A_α_* is the area of cell 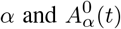 is the preferred area at time *t* (the preferred area will be related with the apicobasal nuclear position in this 2D model, for more details see the following section). The second term describes cell-cell adhesion/tension energy. It introduces the free energy associated with bonds between each cell and its neighbours, where Λ_*ij*_ is a constant and *l_ij_* denotes the length of the junction linking vertices *i* and *j*. When Λ_*ij*_ is negative, cell boundaries tend to expand; when it is positive the edges tend to shrink. The sum of 〈*ij*〉 is over all bonds. The third term describes the contractility of the cell perimeter *L_α_* by a positive coefficient Γ_*α*_, when it is small, contractile forces are small compared to those from area elasticity.

We assume all parameters are the same in each cell or edge, so *K_α_* = *K*, Λ_*ij*_ = Λ, and Γ_*α*_ = Γ.

### Vertex model implementation

The model is implemented by a custom Python code (available in Bitbucket) using the Euler method to solve the equation of movement for each vertex, Eqn (3). We nondimensionalise in time and space by taking as unit of time 460s and using area of 23*μm*^2^. The units of force are arbitrary. The tissue is initialized as an hexagonal mesh of 10 by 10 cells or 15 by 15 cells for simulations with pMn domain. The initial tissue is allowed to evolve for 30h (biological time) to generate a vertex distribution close to steady state. At this point, the time is reset and the vertex distribution is taken as the starting point for the simulations described in this study.

To accommodate topological transitions in the simulations, we introduced the possibility of T1 transitions. During a T1 transition an edge, below 3% of the average edge length of the tissue, is eliminated and a new edge of length *l_new_* expands perpendicular to the old edge (values given in Table 2). If the rearrangement results in the formation of a two-sided cell, the cell is removed from the epithelium.

Division occurs when the cell cycle has been completed and the volume of the cell exceeds a critical value, *A_c_*. In the division process, a new edge is introduced dividing the cell in two and creating two new vertices at the ends of the new edge. The location of the first new vertex is chosen as the midpoint of a randomly selected edge of the dividing cell with probability proportional to the edge length. The other vertex is the midpoint of the opposite edge, that is the edge at the halfway position of the cell from the selected edge. The newly generated sister cells then commence the next cell cycle.

In order to define the frequency of cell divisions in the model, we defined the proliferation rate λ as *d*/(*Ñ* Δ*t*), where *d* is the number of division events in a small time interval Δ*t*, and *Ñ* is the average number of cells in the tissue during Δ*t*. For a proliferating tissue where differentiation does not occur, this estimate of λ is equivalent to the effective rate of tissue growth *k* = ln(*N_C_* (Δ*t*)/*N_C_*)/Δ*t*, where Δ*t* is a time interval, *N_C_* is the number of cells in the tissue at the start of the interval and *N_C_* (Δ*t*) the number of cells at the end of the interval. To match the experimental data (Kicheva et al. (2014), Table 1), in the simulations we aimed to obtain a proliferation rate of 0.05*h*^−1^. For a proliferating tissue without differentiation λ = ln(2)/*t_T_* where *t_T_* is the total cell cycle time. Thus, λ = 0.05*h*^−1^ corresponds to an average cell cycle length of 13*h*, which is 10^5^ simulation time steps (see Table 1 and 2). In tissues with high levels of differentiation, however, the proliferation rate is an effective rate because at any one time a fraction of cells present in the tissue will not further divide. Thus, in a tissue with a differentiation rate of 0.1*h*^−1^ and effective proliferation rate of 0.05*h*^−1^ the cell cycle time of the dividing cells is on average shorter and corresponds to 10.4h. Note that the estimates of cell cycle duration given in Table 1 are based on fractions of dividing cells from fixed images and are therefore effective measurements.

Number neighbour, cell area, and cell perimeter distributions in simulations were compared with output using code from (Smith et al. (2011)).

#### Growth of the tissue

We model the neural tube as a torus with two radii *R* and *H*. The torus can grow in both radial directions. The growth in *R* is resisted by a drag force of magnitude 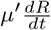 per cell. The forces are balanced so drag forces are of the same total magnitude, as the other forces.

Thus, its growth is determined by a balance between the potential forces and drag:

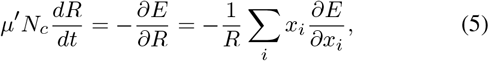

where *x_i_* = *Rθ_i_* is the coordinate that we use for the ith vertex in the direction of dorsoventral (see SM I).

Equivalently, we calculate growth in the perpendicular direction, H, using the drag coefficient, *μ*″.

The tissue aspect ratio (AP/DV) was measured using the whole tissue AP length and DV length.

### Statistical Analysis

Statistical analysis details are documented in figure legends and captions. Kolmogorov-Smirnov tests were used to fit the parameters in Figs 3A, 3B, S.2 and, S.1A, minimising the maximum distance between the experimental and simulated empirical cumulative distribution function of the different observables.

## Acknowledgements

We thank Guillaume Salbreux and Nic Tapon, as well as members of the lab, for constructive comments. We thank Ruth Baker for sharing Aaron Smith’s code (Smith et al. (2011)), with which we tested some of our simulations. This work was supported by: the Francis Crick Institute, which receives its core funding from Cancer Research UK (FC001051), the UK Medical Research Council (FC001051), and Wellcome (FC001051); funding from Wellcome [WT098325MA and WT098326MA] (PG,RP-C, JB and KMP); the European Research Council under European Union (EU) Horizon 2020 research and innovation program grant 680037 (MZ and AK) and 742138 (JB). RP-C acknowledges UCL Mathematics Clifford Fellowship and MZ the National Science Center, Poland (2017/26/D/NZ2/00454).

