## Supplemental material for "Neuronal differentiation affects tissue mechanics and progenitor arrangement in the vertebrate neuroepithelium"

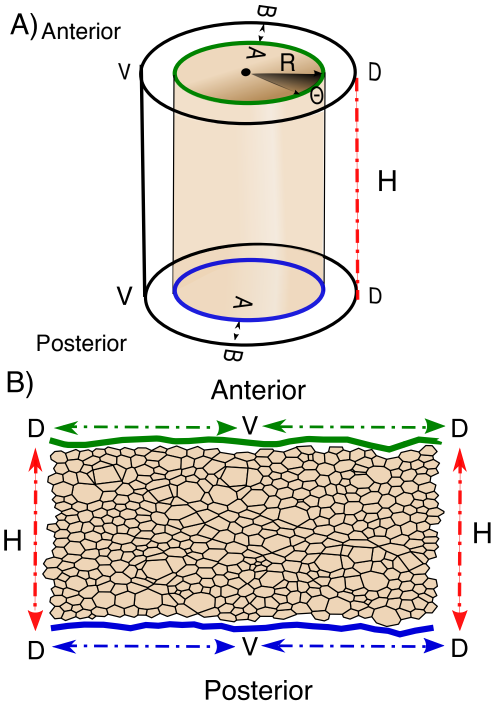


FIG. SM.1. Cylindrical representation where $R$ is the radius, $\Theta$ is the angle and $H$ is height of the cylinder and the $z$-axis is along this length.

### I. CYLINDRICAL REPRESENTATION

The domain of the vertex model is a cylinder, the radius, $R$, and height, $H$, of which can change in time. A natural set of coordinates for the vertex $i$ within the cylinder are $(\Theta_i, \tilde{z}_i)$, where $\Theta_i$ are cylindrical polar angles and $\tilde{z}_i = z_i/H$, where $z_i$ is the coordinate in the direction along the length of the cylinder, see Fig. SM.1. Note that in simulations we actually impose periodic boundary conditions also in the direction along the length of the cylinder. The movement of vertices is determined by an energy function, $E$.

We assume overdamped movement, with the drag coefficient for vertices within the cylinder (whilst the cylinder size is fixed) being $\mu$. Therefore:

$$\mu R \frac{d\theta_i}{dt} = -\frac{1}{R}\frac{\partial E}{\partial \theta_i}, \qquad \text{(SM.1)}$$

$$\mu H \frac{d\tilde{z}_i}{dt} = -\frac{1}{H}\frac{\partial E}{\partial \tilde{z}_i}. \qquad \text{(SM.2)}$$

In addition the cylinder itself can grow. We assume that cells (whose average areas are constant) experience some drag proportional to their speed, both radial and tangential (frictional) as the outside surface of the cylinder expands. Radially the speed of each cell is $\frac{dR}{dt}$ and along the length of the cylinder, the velocities are on average $\frac{1}{2}\frac{dH}{dt}$ relative to a fixed point at one end of the cylinder [1]. Let us define the relevant drag coefficients to be $\mu'$ radially and $2\mu''$ along the length of the cylinders. The equation for the radius and the height are determined by the balance between the radial component (the height component) of the potential forces and the accumulated drag on all of the cells:

$$\mu' N_c \frac{dR}{dt} = -\frac{\partial E}{\partial R}, \mu'' N_c \frac{dH}{dt} = -\frac{\partial E}{\partial H}, \qquad \text{(SM.3)}$$

where $N_c$ is the number of vertices. We note that the three drag coefficients $\mu$, $\mu'$ and $2\mu''$ are likely to be different. We also note when a cell is added its energy is added to $E$.

The coordinates that we actually use in the vertex model code are given by $(x_i, z_i)$, where $x_i = R\theta_i$ and $z_i = H\tilde{z}_i$, which are effectively rectangular coordinates in a folded out cylinder. The equation for the radius, thus becomes:

$$N_c \mu' \frac{dR}{dt} = -\sum_i \frac{\partial E}{\partial x_i}\frac{\partial x_i}{\partial R} = -\frac{1}{R}\sum_i x_i \frac{\partial E}{\partial x_i}. \qquad \text{(SM.4)}$$

Therefore

$$\frac{dx_i}{dt} = -\frac{1}{\mu}\frac{\partial E}{\partial x_i} - \frac{1}{\mu'}\frac{x_i}{N_c R^2}\sum_j x_j \frac{\partial E}{\partial x_j}. \qquad \text{(SM.5)}$$

Similarly,

$$\frac{dz_i}{dt} = -\frac{1}{\mu}\frac{\partial E}{\partial z_i} - \frac{1}{\mu''}\frac{z_i}{N_c H^2}\sum_j z_j \frac{\partial E}{\partial z_j}. \qquad \text{(SM.6)}$$

In the code, we update $x_i$ and $z_i$ first according to equation (SM.5) and (SM.6), (i.e. in time step $\Delta t$ we add $R\Delta\theta_i$ and $H\Delta\tilde{z}_i$), then we multiply by a factor $\frac{R+\Delta R}{R}$ and $\frac{H+\Delta H}{H}$ respectively, where we call $\frac{\Delta R}{R}$ and $\frac{\Delta H}{H}$ "expansion" and these are given by $-\frac{\Delta t}{N_c \mu' R^2}\sum_j x_j \frac{\partial E}{\partial x_j}$ and $-\frac{\Delta t}{N_c \mu'' H^2}\sum_j z_j \frac{\partial E}{\partial z_j}$.

[1] The magnitude of the overall force is the same if we consider the middle of the cylinder as being stationary.

### II. BIOPHYSICAL PARAMETER SPACE

We study the effect of the biophysical parameters on the cell dynamics. The energy (Eq. 4) is determined by the vertex positions and parameters $K$, $\Lambda$, and $\Gamma$. In Magno et al. [2015] find parameter regions that yield different behaviours depending on the mechanical parameter.

The energy $E$ is given by:

$$E = \frac{K}{2}(A - A^0)^2 + \Lambda L + \frac{\Gamma}{2}L^2. \quad \text{(SM.7)}$$

The interfacial tension $\gamma$ and pressure $\Pi$ are given by

$$\gamma = \frac{\partial E}{\partial L} = \Lambda + \Gamma L, \quad \text{(SM.8)}$$

$$\Pi = -\frac{\partial E}{\partial A} = -K(A - A^0). \quad \text{(SM.9)}$$

In order for a cell to minimise its perimeter and therefore take a regular shape, we require the interfacial tension to be positive, hence

$$\gamma = \Lambda + \Gamma L > 0. \quad \text{(SM.10)}$$

This should be true for the equilibrium cell size.

Suppose a cell has a fixed shape and denote the square root of its area by $l$ and its shape index by $s$. The equilibrium size is determined by setting $\frac{\partial E}{\partial l} = 0$:

$$\frac{\partial E}{\partial l} = \gamma \frac{dL}{dl} - \Pi \frac{dA}{dl} = 0, \quad \text{(SM.11)}$$

thus at equilibrium $\gamma s = 2\Pi l$, and therefore

$$2Kl^3 = -\Lambda s + (2KA^0 - \Gamma s^2)l. \quad \text{(SM.12)}$$

We note that when $l = 0$, $\frac{\partial E}{\partial l} = \Lambda$, so the state $l = 0$ is stable if and only if $\Lambda > 0$. Eq. SM.12 has a single positive solution corresponding to an energy minimum if $\Lambda < 0$ (meaning cells converge to a single equilibrium area). If $\Lambda > 0$, Eq. SM.12 can either have zero positive solutions (in which case all cells collapse to zero area) or two positive solutions. In the latter case, the larger positive solution will correspond to an energy minimum and the small positive solution to an energy maximum. Thus cells with area greater than the unstable value will expand their area towards a fixed positive value, while the area of small cells will collapse to zero.

We note that the condition for the state $l = 0$ to be unstable is independent of the target area. In our model, the target area varies slowly compared to the movement of vertices and thus we expect this property to hold for the same parameter values in the model with varying target area too. The condition for regularly shaped cells does depend on the target area, since when interfacial tension is zero, the minimum energy will be attained when cell area is equal to target area. With other parameters fixed, as the target area increases, the interfacial tension will also increase and so cells will tend to be more regular. Thus it is possible that small cells will be irregularly shaped, but larger ones will be regularly shaped. This is because for small cells the adhesive force will dominate the force from the actomyosin ring, whereas for larger cells, the opposite will be true.

The final condition that we need to determine is the condition for there to be two positive steady states instead of zero when $\Lambda > 0$. There are two positive states if and only if

$$A^0 > \frac{\Gamma s^2}{2K} + 3\left(\frac{\Lambda s}{4K}\right)^{2/3}. \quad \text{(SM.13)}$$

This condition depends on our variable target area. What really matters, however, is whether cells are likely to grow at any size that they attain (i.e. it does not matter if we are in the region with two stable areas if the cells are so small when they are born that they shrink to zero). A necessary condition for cells to always grow is that $\frac{\partial E}{\partial l}\Big|_{l=\sqrt{A_c/2}} < 0$ with the target area given by its value at the start of the cell cycle. A sufficient condition is that $\frac{\partial E}{\partial l}\Big|_{l=\sqrt{A_c/2}} < 0$ with the target area given by its minimal value. Now

$$\left.\frac{\partial E}{\partial l}\right|_{l=\sqrt{A_c}/2} = \Lambda s + \Gamma s^2\sqrt{A_c/2} + 2K\sqrt{A_c/2}(A_c/2 - A^0). \quad \text{(SM.14)}$$

The cells will thus grow if

$$2K\sqrt{A_c/2}(A^0 - A_c/2) > \Lambda s + \Gamma s^2\sqrt{A_c/2}. \quad \text{(SM.15)}$$

Let us use the notation of Farhadifar et al. [2007], with the $A^0$ that they use to create the nondimensional parameters given by its mean value, $\langle A^0\rangle$ during the cell cycle in our model. The nondimensional parameters are $\bar{\Lambda} = \frac{2\Lambda}{K\langle A^0\rangle^{3/2}}$ and $\frac{\Gamma}{K\langle A^0\rangle}$. Let $A_c/(2\langle A^0\rangle) = c$ and the value of $A^0$ in Eq. SM.15 be $\beta\langle A^0\rangle$. Then cells will grow if

$$\bar{\Lambda}s/2 + \bar{\Gamma}s^2\sqrt{c} < 2\sqrt{c}(\beta - c). \quad \text{(SM.16)}$$

If the model of target area variation with the cell cycle and the critical area for cell division are fixed, then, for a given shape index (e.g. that of a regular hexagon), this forms a diagonal region in the phase space.

We note that equations SM.13 and SM.15 imply that cells with lower shape indices (i.e. more regular and with larger number of sides) are more likely to be stable.

In Fig. SM.2, we show the phase diagram in the space of the parameters $\bar{\Lambda}$ and $\bar{\Gamma}$. Region I is where cells are expected to have negative interfacial tensions and so have irregular shapes since they do not want to minimise their perimeters. The boundary of this domain is at $\Lambda + \Gamma s\sqrt{A^0} = 0$, so $\bar{\Lambda} + 2\bar{\Gamma}s = 0$. We plot this for hexagonal cells ($s = 2 \times 3^{1/4}2^{1/2}$). In Region II,

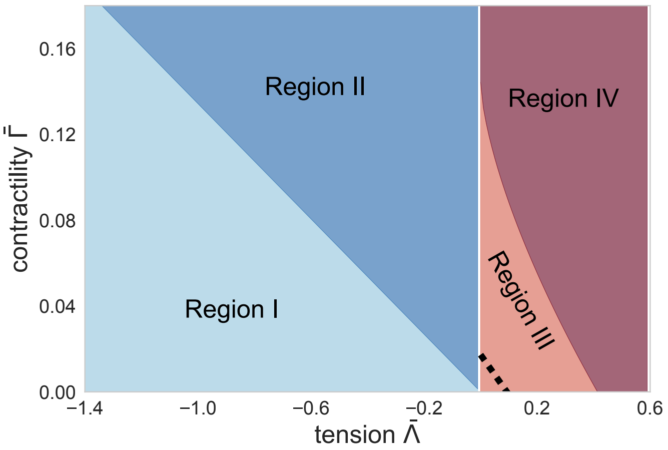


FIG. SM.2. Diagram phase for biophysical parameter space for hexagonal cell in term of $\bar{\Lambda}$ and $\bar{\Gamma}$.

cells take regular shapes and the only stable size is positive. The other boundary of this domain is at $\Lambda = 0$ or $\bar{\Lambda} = 0$. In Region III, cells take regular shapes and small cells collapse, whilst larger ones grow to an equilibrium size. In Region IV, all cells collapse. The boundary between these regions is given by $A^0 = \frac{\Gamma s^2}{2K} + 3\left(\frac{\Lambda s}{4K}\right)^{2/3}$ or $2 = s^2\bar{\Gamma} + \frac{3}{2}(s\bar{\Lambda})^{2/3}$. We plot this for hexagonal cells. We also show as the dotted line, a line below which all cells should grow (Eq. SM.16), with $\beta$ given by the ratio of the minimal value of $A^0$ and its mean value.

### III. TISSUE ASPECT RATIO DEPENDS ON EXPANSION RATE, THE VALUE OF $\bar{\Gamma}$ AND ON HOW EFFICIENTLY CELLS REARRANGE

To gain insight into the anisotropic growth of the tissue, we consider a slightly simpler system. We assume all cells are identical and we assume that $\Lambda/\Gamma << 1$, so that the target perimeter is negligible. Let the dimensions of the neural tube be $R(t)$ and $H(t)$ and the number of cells be $N_c(t)$. Therefore $N_c A = RH$. For ease of analysis, we assume that cells are rectangular. In the direction of $R$, there are $n_1$ cells of length $l_1$, with $R = n_1 l_1$ and similarly in the other direction there are $n_2$ cells of length $l_2$. Therefore, $R/H = (n_1\, l_1)/(n_2\, l_2)$. We start with an equal number of square cell in each direction. We consider two extreme cases. In the first case, intercalation can take place rapidly, allowing the number of cells in each direction to change. In this case, any change in the tissue shape and the dissipation of drag forces will occur predominantly by changing the number of cells in each direction, rather than by altering cell shape. In such a scenario, the shape of individual cells will be relatively unaffected by anisotropic forces and will be square, hence $n_1/n_2 = R/H$. In the second case, intercalation does not take place, in which case as the aspect ratio of the whole tissue changes, the cells change shape accordingly. In this case, the number of cells in each direction does not change, since cell division is unorientated and intercalation impossible, hence $n_1 = n_2$ and $l_1/l_2 = R/H$.

The expansion of the tissue in dimensions $R$ and $H$ and perimeter $P = 2(l_1 + l_2)$, is given by SM.3. In turn,

$$\begin{aligned}\partial E/\partial R &= N_c K(A - A_0)\partial A/\partial R + N_c \Gamma P \partial P/\partial R \\ &= HK(A - A_0) + 2N_c\Gamma P/n_1 \\ &= HK(RH/N_c - A_0) + 4n_2\Gamma(l_1 + l_2) \\ &= H[K(RH/N_c - A_0) + 4\Gamma(1 + l_1/l_2)].\end{aligned} \tag{SM.17}$$

Similarly,

$$\partial E/\partial H = R\left[K(RH/N_c - A_0) + 4\Gamma(1 + l_2/l_1)\right]. \tag{SM.18}$$

In the first case $\mu''/\mu'\, dR/dH = H/R$ and the change in $R^2$ will be $\mu''/\mu'$ times the change in $H^2$, so the aspect ratio will tend to $\sqrt{\frac{\mu''}{\mu'}}$.

In the second case,

$$\mu' N_c \frac{dR}{dt} = H[K(A_0 - RH/N_c) - 4\Gamma(1 + R/H)] \tag{SM.19}$$

$$\mu'' N_c \frac{dH}{dt} = R[K(A_0 - RH/N_c) - 4\Gamma(1 + H/R)]. \tag{SM.20}$$

We expect that in the long term the growth will equilibrate so that the average area of cells will tend to a constant, i.e. $RH/N_c$ will tend to a constant. This constant will depend on the rate of proliferation. The faster the rate of proliferation (actually the exponential growth rate of $N_c$) compared to the movement of vertices, the smaller this area will be [2]. We let $B = K\,A_0 - 4\Gamma - K\,H\,R/N_c = K\,A_0 - 4\Gamma - KA$, so the faster $N_c$ increases compared to the rate of movement of vertices , the smaller $A = H\,R/N$ will be, so the larger B will be. We obtain

$$\mu' N_c \frac{dR}{dt} = BH - 4\Gamma R \tag{SM.21}$$

$$\mu'' N_c \frac{dH}{dt} = BR - 4\Gamma H. \tag{SM.22}$$

We note that the tissue will only grow if $B > 4\Gamma$, which implies $\bar{\Gamma} < 1/8$. Letting $r = R/H$, we get

$$N_c dr/dt = \frac{B}{\mu'} + 4\Gamma r\left[\frac{1}{\mu''} - \frac{1}{\mu'}\right] - r^2\frac{B}{\mu''}. \tag{SM.23}$$

This means that as $t \to \infty$, $r \to \frac{2\Gamma}{B}\left(1 - \frac{\mu''}{\mu'}\right) + \sqrt{\frac{4\Gamma^2}{B^2}\left(1 - \frac{\mu''}{\mu'}\right)^2 + \frac{\mu''}{\mu'}}$.

[2] Cell division does not require movement of vertices, just the addition of two new vertices.

In the limit of very slow growth with $B \approx 4\Gamma$, this is approximately 1, so that very slow growth is almost isotropic. For faster proliferation rates B is higher, hence r diverges from 1 and the tissue grows anisotropically. In the limit, $\Gamma/B \to 0$, the aspect ratio tends to $\sqrt{\frac{\mu''}{\mu'}}$. Increasing the proliferation rate, decreases $\Gamma/B$ from $\frac{1}{4}$ to $\frac{\bar{\Gamma}}{1-4\bar{\Gamma}}$. Thus for a fixed value of $\bar{\Gamma}$, even extremely rapidly growing tissues will not achieve aspect ratios as extreme as $\sqrt{\frac{\mu''}{\mu'}}$. The most extreme aspect ratios are achieved when the actomyosin ring exerts a negligible force or when tissue rearrangement is extremely efficient.

**Farhadifar, R., Röper, J.-C., Aigouy, B., Eaton, S. and Jülicher, F.** (2007). The influence of cell mechanics, cell-cell interactions, and proliferation on epithelial packing. *Current Biology* **17**, 2095 – 2104. ISSN 0960-9822.

**Magno, R., Grieneisen, V. A. and Marée, A. F.** (2015). The biophysical nature of cells: potential cell behaviours revealed by analytical and computational studies of cell surface mechanics. *BMC Biophysics* **8**, 8. ISSN 2046-1682.
