## Supplemental Figure 1 for "Neuronal differentiation affects tissue mechanics and progenitor arrangement in the vertebrate neuroepithelium"

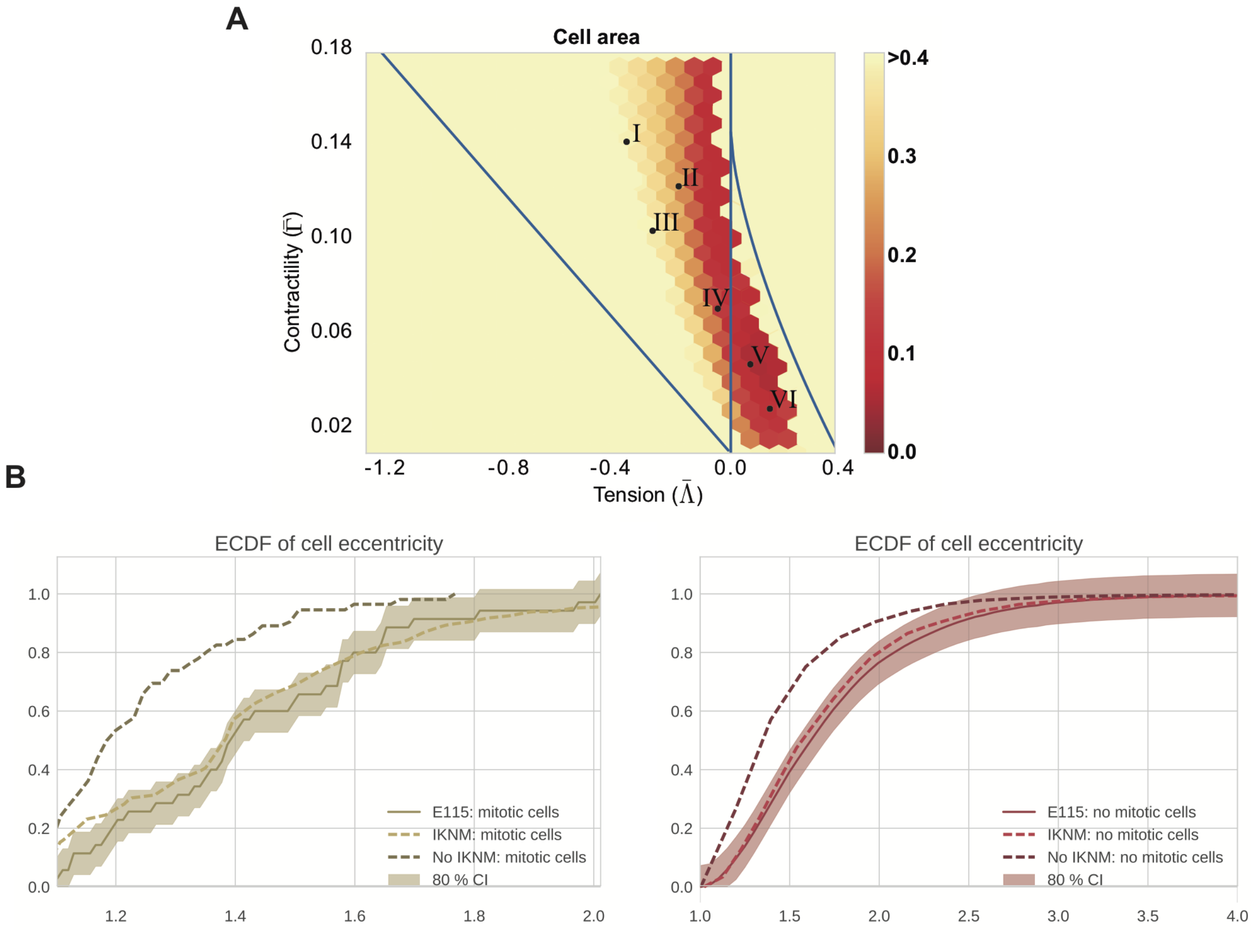


Figure S.1: A) Map of the average of maximum distance between the experimental and simulated empirical cumulative distribution function of cell area distribution for a model without IKNM (constant target area). Experimental data corresponds to E11.5 embryos and simulations correspond to 10 simulations per point in the parameter space. Roman numeral marked points indicate the mechanical parameters selected, $(\bar{\Lambda}, \bar{\Gamma})$ in Fig. 3A. B) Empirical cumulative distribution function (ECDF) of cell angle orientation and eccentricity in simulations with and without IKNM using mechanical parameter V in Fig. 3A compared to E11.5 experimental data of mitotic and non-mitotic cells.
