## Supplemental Figure 2 for "Neuronal differentiation affects tissue mechanics and progenitor arrangement in the vertebrate neuroepithelium"

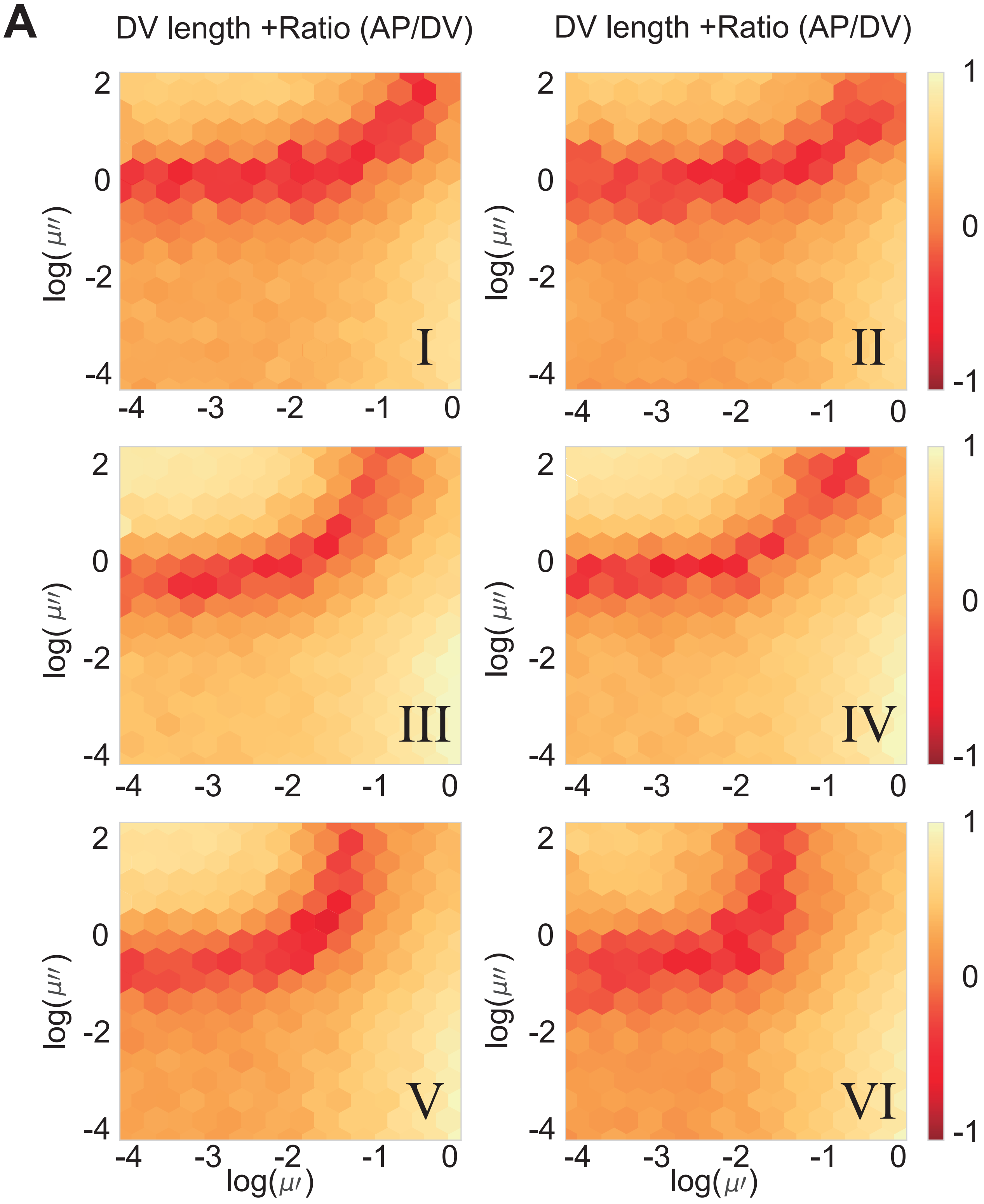


Figure S.2: A) Heatmap indicating the difference between simulation and experimental tissues for different values of the drag coefficients $\mu'$ and $\mu''$. Differences between experiments and model were evaluated by comparing the sum of DV growth of the tissue and tissue aspect ratio (AP/DV) after 48h of simulation time. Colour map shows the average difference (log scale) from 10 different simulations per point with the experimental data from E11.5 embryos. Other parameters are indicated in Table 2. Roman numerals indicate mechanical parameters used $(\bar{\Lambda}, \bar{\Gamma})$ in Fig. 3A.
