## Supplemental Figure 3 for "Neuronal differentiation affects tissue mechanics and progenitor arrangement in the vertebrate neuroepithelium"

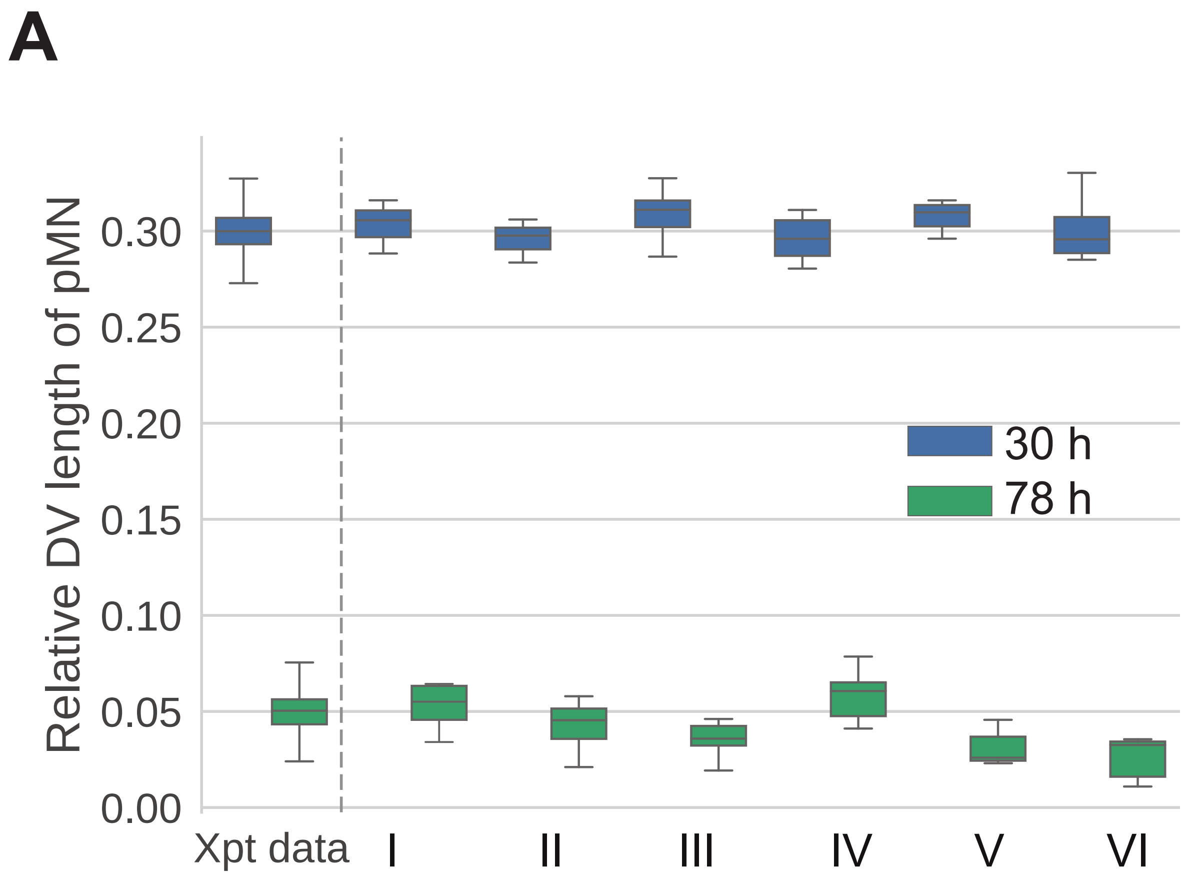


Figure S.3: A) DV length of the pMN domain relative to the total DV length of the tissue at the moment pMN is introduced to simulations (blue) and after 48 hours (green). Differentiation rate in pMN domain $0.1h^{-1}$. Other parameters are indicated in Table 2. These time points correspond to experimental samples (Xpt data) from E9.5 and E11.5 mouse embryos, respectively. The DV length at these developmental stages was measured in (Kicheva et al. (2014)).
