## Supplemental Figure 4 for "Neuronal differentiation affects tissue mechanics and progenitor arrangement in the vertebrate neuroepithelium"

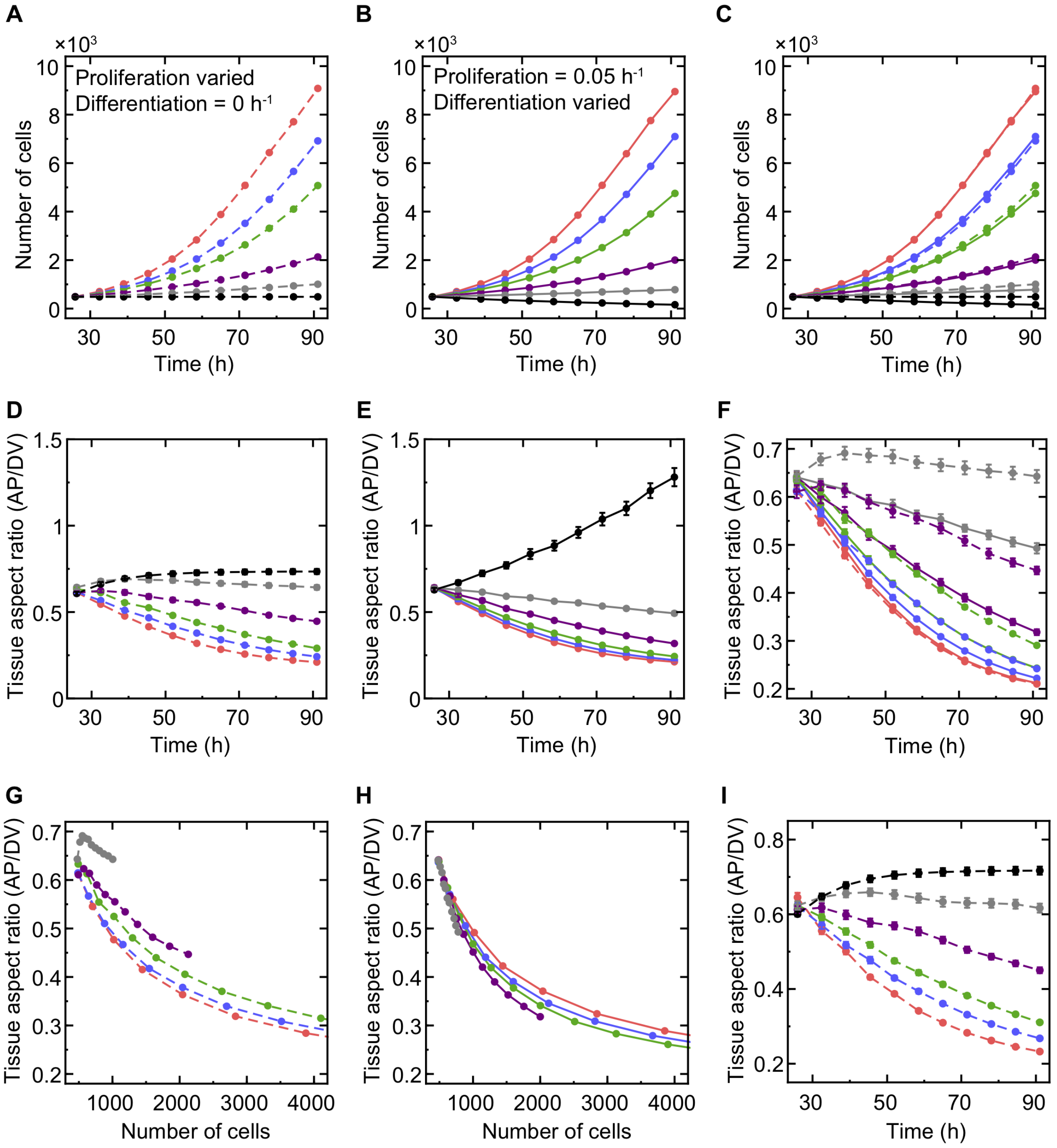


Figure S.4: A) Number of cells in simulated tissues as a function of time for different proliferation rates and no differentiation. The proliferation rates from top to bottom: $0.05h^{-1}$ (red), $0.04h^{-1}$ (blue), $0.03h^{-1}$ (green), $0.02h^{-1}$ (purple), $0.01h^{-1}$ (grey), $0h^{-1}$ (black). In A-I: the simulation starts with an initial period of ($t < 26h$) during which the proliferation rate is $0.05h^{-1}$ and there is no differentiation. Afterwards proliferation or differentiation is altered as indicated. In A-H parameter regime V is used. B) The number of cells as a function of time for different differentiation rates and fixed proliferation rate = $0.05h^{-1}$. The differentiation rate is $0h^{-1}$ (red), $0.01h^{-1}$ (blue), $0.02h^{-1}$ (green), $0.035h^{-1}$ (purple), $0.05h^{-1}$ (grey), $0.075h^{-1}$ (black). The parameters were chosen so that the net growth rate in B is similar to that in A for curves of the same colour (except black). C) Overlaid data from A and B. D) Aspect ratio of the tissue AP to DV extension for varied proliferation and no differentiation (same as main Fig. 4F, parameters as in A). E) As D but for varied differentiation and fixed proliferation (parameters as in B). F) Overlaid data from D and E. Only simulations with positive net growth rate are plotted (no black trajectories). G, H) The tissue aspect ratio (AP/DV) from D and E as a function of cell number. Only conditions with positive growth rate are shown. I) The tissue aspect ratio (AP/DV) for varied proliferation and no differentiation in parameter regime VI. The proliferation rates and colour convention as in (A). Data A-J was averaged over 12 independent simulations for each condition. The reported errors are SEM. If error bars are not visible the SEM was very small (<2%).
