## Supplemental Figure 5 for "Neuronal differentiation affects tissue mechanics and progenitor arrangement in the vertebrate neuroepithelium"

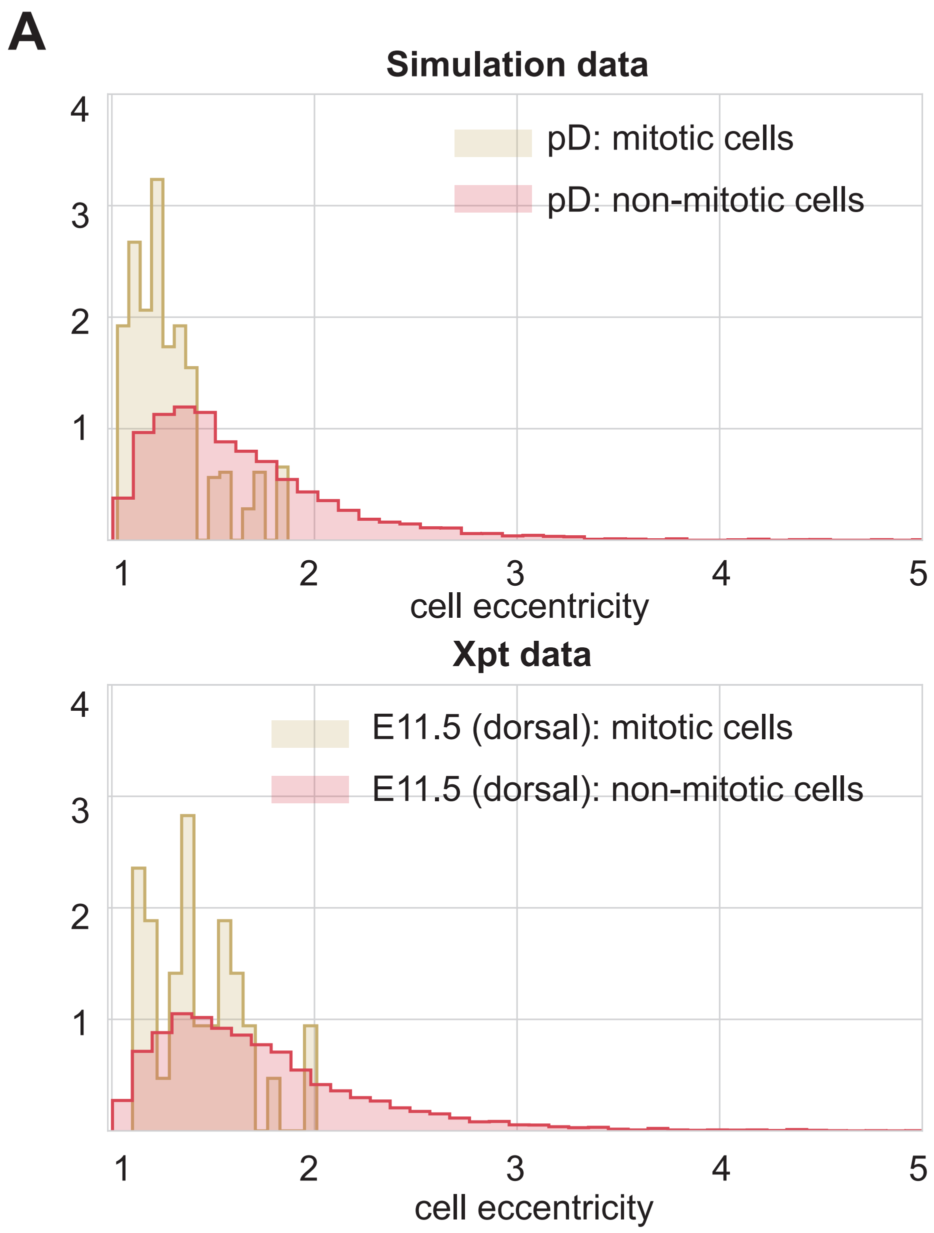


Figure S.5: A) Histogram of eccentricities of cells close to mitosis (brown) and non-mitotic (red) cells in simulations and experimental data in dorsal domain. In the experimental data, the 2.5% of cells with largest area were considered mitotic.
